# Mitochondria-derived nuclear ATP surge protects against confinement-induced proliferation defects

**DOI:** 10.1101/2023.12.20.572417

**Authors:** Ritobrata Ghose, Fabio Pezzano, Savvas Kourtis, Ilir Sheraj, Shubhamay Das, Antoni Gañez Zapater, Upamanyu Ghose, Lorena Espinar, Katja Parapatics, Valeria Venturini, André C Müller, Verena Ruprecht, Sara Sdelci

## Abstract

The physical microenvironment regulates cell behaviour. However, whether physical confinement rewires the subcellular localisation of organelles and affect metabolism is unknown. Proteomics analysis revealed that cellular confinement induces a strong enrichment of mitochondrial proteins within the nuclear compartment. High-resolution microscopy confirmed that mechanical cell confinement leads to a rapid re-localisation of mitochondria to the nuclear periphery. This nuclear-mitochondrial proximity is mediated by an endoplasmic reticulum-based net that entraps the mitochondria in an actin-dependent manner. Functionally, the mitochondrial proximity results in a nuclear ATP surge, which can be reverted by the pharmacological inhibition of mitochondrial ATP production or via actin depolymerisation. Inhibition of the confinement-derived nuclear ATP surge reveals long-term effects on cell fitness which arise from alterations of chromatin states, delayed DNA damage repair, and impaired cell cycle progression. Together, our data describe a confinement-induced metabolic adaptation that is required to enable prompt DNA damage repair and cell cycle progression by allowing chromatin state transitions.

## INTRODUCTION

Actively proliferating cells, such as cancer cells within the tumour mass or embryonic cells during development, are subjected to physically confining environments that expose them to acute mechanical stress and can alter cytoskeleton organisation^1–3^, nuclear structure^4^, and chromatin organisation^5^. However, despite the presence of mechanical confinement stress in various biological processes, as well as the accumulation of DNA damage and mitotic defects^6–8^, cell proliferation persists, suggesting various unexplored adaptation mechanisms in play. Mechanical signals from the microenvironment may trigger both physical and functional adaptive processes that promote cell fitness and survival. Mechanical remodelling of intracellular organelles has been shown for the nucleus^4^ and mitochondria^9,10^. Nevertheless, it remains unknown whether these processes are required for rapid metabolic rewiring or are driven by an active reorganisation of organelle subcellular redistributions and crosstalk.

Here, we studied how the localisation of cellular organelles and molecular components changes following acute mechanical confinement and how they can trigger rapid metabolic cell changes relevant for mechano-adaptive cell responses involved in stress resistance and cell fitness. We performed a proteomics-based subcellular fractionation of acutely confined cells, which allowed us to identify the redistribution of proteins within the nucleus and cytoplasmic compartment. We found that upon mechanical cell deformation in confinement, mitochondria accumulate within the nuclear area and are trapped within an immobilised ER network in an actin-dependent manner. This leads to a multi-organelle architectural reorganisation, involving mitochondrial accumulation at the nuclear periphery that deform the nucleus and lead to characteristic nucleus shape changes. The close proximity between mitochondria and the nucleus is further responsible for mechanical stress induced metabolic changes within the nucleus in the form of increased ATP levels of mitochondrial origin. We functionally show that these elevated nuclear ATP levels are required to allow changes in chromatin accessibility which are required for an efficient DNA damage repair signalling and the proper progression of the S-phase of the cell cycle, safeguarding cell proliferation via a metabolic cell mechano-adaptation mechanism during acute confinement stress.

## RESULTS

### Cell confinement induces a mitochondrial enrichment at the nucleus

To explore cellular adaptation mechanisms to mechanical stress and study the contribution of specific subcellular compartments, we performed a subcellular fractionation of HeLa cells acutely confined for 15 minutes. HeLa cells are considered as confined when deformed to a height of 3 μm, resulting in cytoskeleton reorganisation and bleb formation^11^. To achieve this, we adapted a previously established agarose-based confiner^11^, where an agarose disc applies pressure onto cells, simulating a physically constrained environment (Figure 1 A). Using confocal microscopy of confined HeLa cells expressing a whole-cell green fluorescent protein (GFP) marker, we validated the cell confinement height and mechano-sensitive bleb formation (Figure S1 A) as previously documented^1,11^. To investigate relative changes in the nucleus versus cytoplasmic cell compartments we performed a nucleo-cytoplasmic fractionation under confined conditions. To achieve this, we injected a cytoplasmic lysis buffer through the agarose pad, following an acute 15-minute confinement (Figure 1 A). The lysate was then centrifuged to obtain a nuclear protein fraction and a cytoplasmic protein supernatant. The two fractions were validated using Western Blot analysis where Vinculin, a cytoskeletal protein, and nuclear histone H3 were detected within either the cytoplasmic or nuclear fractions, respectively (Figure S1 B). Unconfined HeLa cells in suspension (henceforth referred to as suspension cells) were lysed to obtain reference cytoplasmic and nuclear fractions.

**Figure 1.**
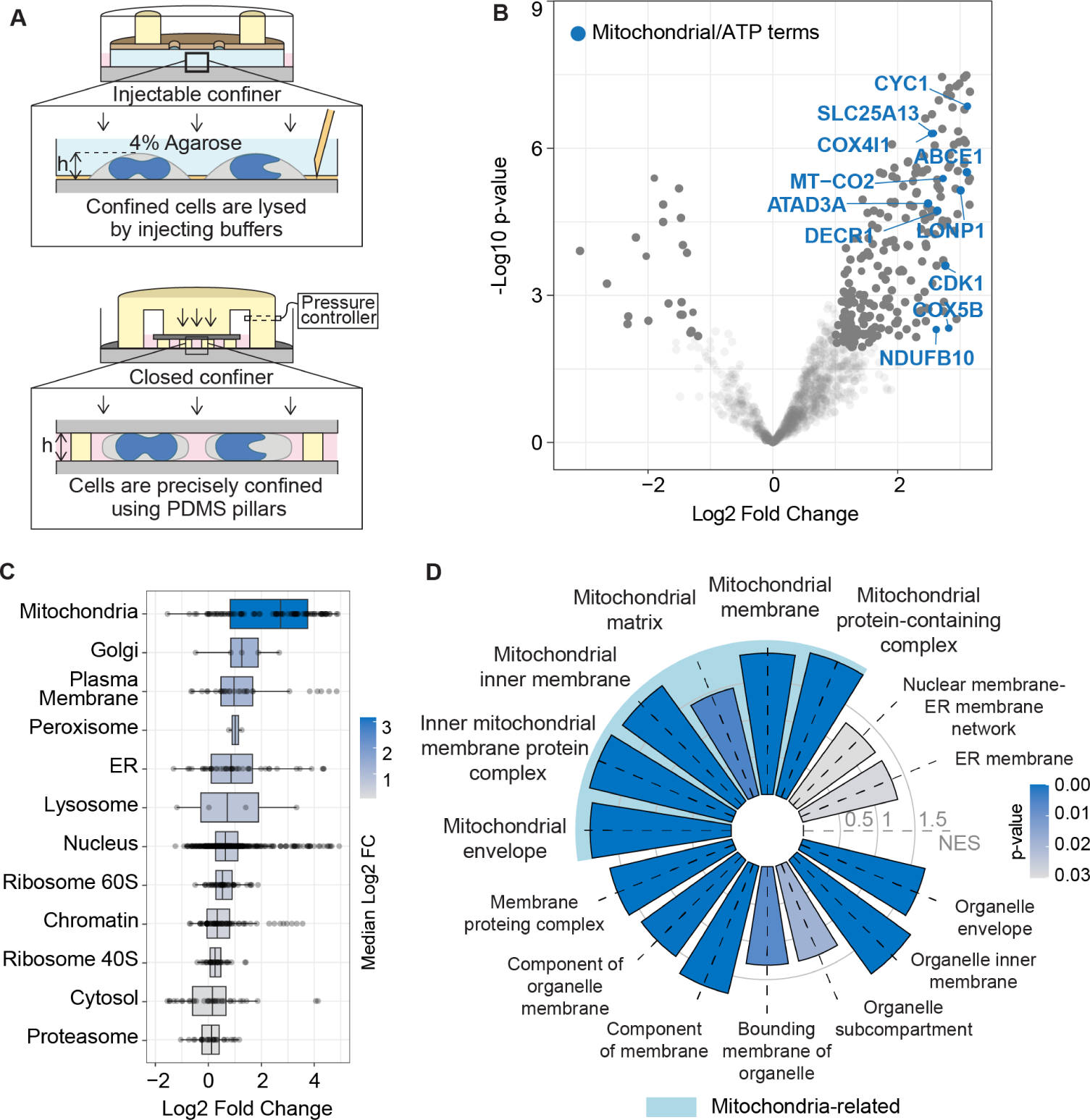
Proteomic analyses reveal mitochondrial enrichment within the nuclear fraction. **(A)** Schematic of the two confinement systems used in this study: [top] Injectable confiner, which is based on a 4% agarose pad for confinement through which reagents can be injected, and [bottom] a pressure controlled closed confinement system using PDMS microspacers. **(B)** Enriched and depleted terms when comparing the nuclear fraction of confined HeLa cells to that of unconfined cells. **(C)** Comparison of the enriched terms from the nuclear fractions in (B), mapped to organelles based on their known localisation from the hyperLOPIT dataset^12^. **(D)** Cellular components – GSEA results performed on the nuclear fractions in (B). Light blue background highlights terms related to mitochondria. Concentric circles represent normalised enrichment scores (NES). See also Figure S1 for quality control of subcellular fractionation and proteomics.

A principal component analysis of mass spectrometry data showed a clear separation between the nucleus and the cytoplasm, as well as between confinement and suspension, suggesting an altered proteomic distribution within these subcellular compartments upon confinement (Figure S1 C). As expected, proteins known to be localised to the nucleus^12^ were enriched in the nuclear fraction of suspension cells (Figure S1 D). Gene set enrichment analysis (GSEA) of these nuclear fraction-associated proteins compared to cytoplasmic fraction-related proteins showed an enrichment of nuclear and chromatin related gene ontologies (GOs) and a depletion of cytoplasm-related GOs in suspension (Figure S1 E) and in confined cells (Figure S1 F-G), validating our approach.

While the comparison of cytoplasmic fractions did not reveal a pronounced protein redistribution (Figure S1 H-I), comparing the two nuclear fractions notably revealed a marked enrichment of mitochondrial proteins in confined nuclei (Figure 1 B), which was much more pronounced than the enrichment of other cellular organelles (Figure 1 C). GSEA of enriched proteins within the nuclear fraction also confirmed this enrichment of the mitochondria (Figure 1 D).

Altogether, our proteomics approach revealed that acute confinement provokes an enrichment of mitochondria in the nuclear fraction.

### Mitochondrial accumulation promotes nuclear shape changes

The presence of mitochondria-related terms within the nuclear fraction of confined cells may be due to a stronger association between mitochondria and the nuclear surface or be a consequence of mitochondria being inside the nucleus. To test which of these hypotheses could explain the mitochondrial enrichment in our proteomic data, we performed high-resolution confocal microscopy of suspension and confined HeLa cells and monitored mitochondria localisation with respect to the nucleus using MitoTracker and Hoechst stains. Suspension and confined cells both showed a strong cytoplasmic MitoTracker signal. However, in stark contrast to suspension cells, where the nuclear region was devoid of mitochondria, confined cells showed a strong enrichment of MitoTracker signal within the nuclear region (Figure 2 A). Three-dimensional (3D) image reconstructions showed that while in suspension cells mitochondria were homogeneously distributed in the cytoplasm, in confined cells mitochondria were enriched within the nuclear periphery, and notably above and below the nucleus (Figure 2 B). Moreover, the mitochondrial signal was mutually exclusive to the Hoechst signal, indicating that they were not physically inside the nucleus. Notably, 3D image reconstruction allowed us to observed characteristic nuclear shape changes in the confined cells, with indentations on the top, bottom, or both the top and bottom of the nucleus, within which mitochondria were found (Figure 2 C). Image analysis of confined HeLa cells, transiently overexpressing the inner nuclear membrane marker LAP2β-GFP, also confirmed that during confinement mitochondria were redistributed and accumulated within nuclear indentations, while remaining at the outer nuclear periphery (Figure 2 D). Line profiles of MitoTracker and Hoechst signal intensities across confined cells clearly showed the signals of mitochondria and the nucleus overlap, while suspension cells showed mutually exclusive signals (Figure 2 E). Together, these observations corroborated the presence of mitochondria-enriched nuclear indentations upon mechanical cell confinement.

**Figure 2.**
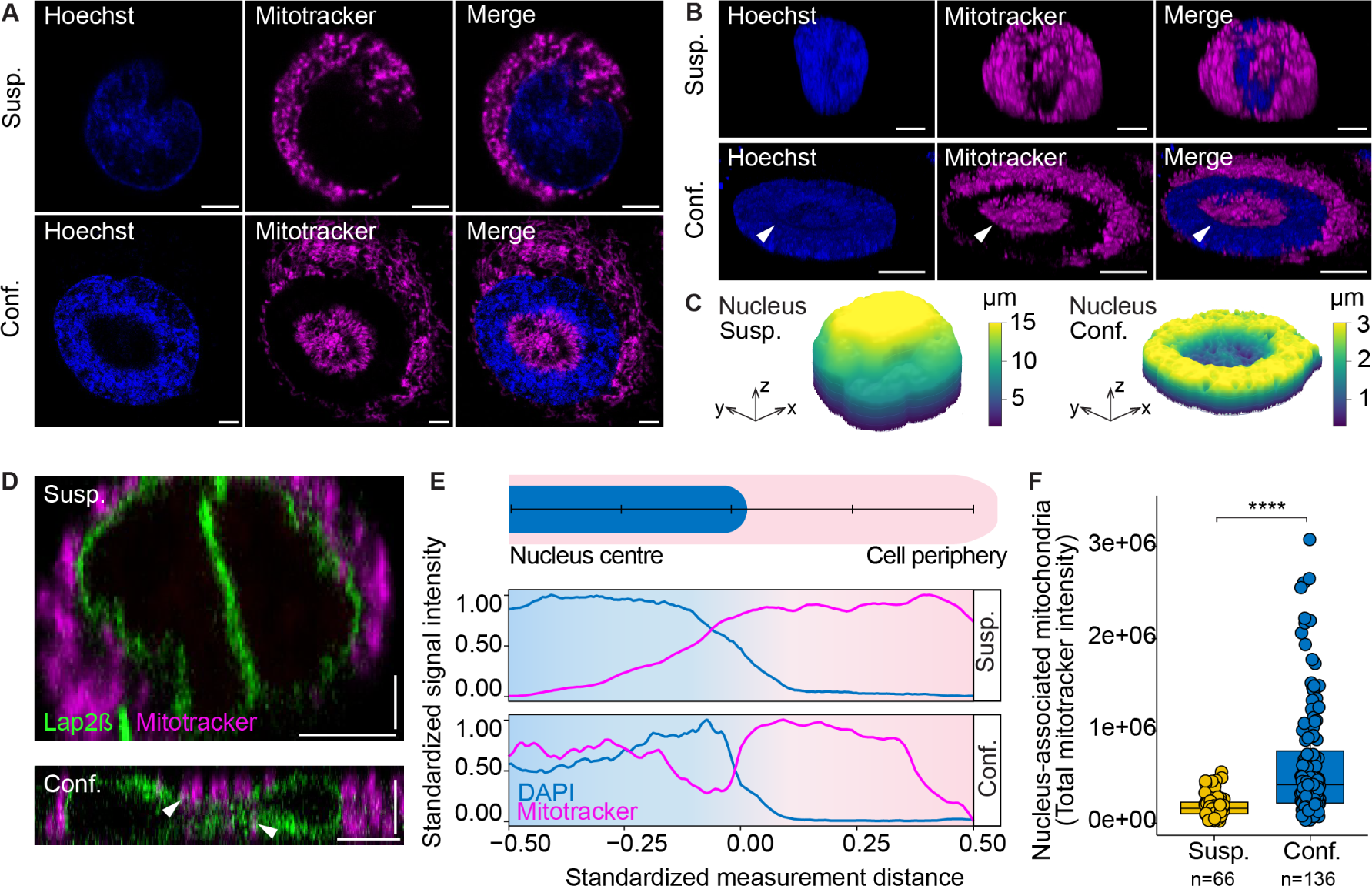
Confinement induces nuclear indentations and accumulation of mitochondria at the nuclear periphery. **(A)** HeLa cells were stained with Hoechst to visualise nuclei and MitoTracker to label mitochondria in control suspension cells and confined cells. **(B)** 3D representation of suspension and confined cells shown in (A). White arrow highlights an indentation within the nuclear surface where mitochondria are present. **(C)** Hoechst signal was used for an in silico reconstruction of the nucleus of the suspension and confined cells shown in (A-B). **(D)** LAP2β-GFP-expressing and MitoTracker-stained HeLa cells were used to visualise the inner nuclear membrane and mitochondrial localisation during confinement. [top] X-Z view of a suspension HeLa cell, and [bottom] X-Z view of a confined HeLa cell. White arrows highlight indentations at the top and bottom of the nucleus within which mitochondrial signal is detected. **(E)** Hoechst and MitoTracker line profiles from the centre of the nucleus to the cell periphery in suspension and confined HeLa cells. **(F)** Nucleus-associated mitochondria (NAM) were quantified as mitochondria situated between the nucleus and 10% of its surrounding area in suspension and confined HeLa cells. All scale bars, except vertical scale bars in (D), represent 5 μm. Vertical scale bars in (D) are 3 μm. Statistics in (F) were performed using the Wilcoxon test (* P<0.05, **** P<0.0001). See also Figure S2 for characterisation of nuclear indentation and NAM, and validation in other cell lines.

Nuclear indentations could be classified into different shapes as “doughnuts”, “beans”, or more complex shapes based on the Hoechst signal and localisation of mitochondria. Doughnuts contained indentations either on the top, bottom, or both the top and bottom of the nucleus, bean nuclei contained indentations on the side, and complex shapes contained multiple indentations, and could not be classified into either doughnuts or beans. In HeLa cells, doughnut shaped nuclei were the most abundant (75%), followed by beans (24%) and complex nuclei (2%) (Figure S2 A). To assess the penetrance of this nucleus-associated-mitochondria (NAM) phenotype within the HeLa cell population, we classified confined cells as ‘NAM-present’ or ‘NAM-absent/low’ and found 84% of the population displaying a strong NAM phenotype (Figure S2 B), indicating that the large majority of cells underwent a nuclear-mitochondrial physical remodelling upon mechanical cell confinement.

We next assessed the presence of this phenotype in different cell lines and applied acute 3 μm confinement to the osteosarcoma cell line U2OS and the triple negative breast cancer cell line MDA-MB-231. With a variable penetrance, both U2OS and MDA-MB-231 cells showed a consistent NAM increase when confined (Figure S2 B), suggesting that the NAM phenotype may be a conserved response to mechanical confinement.

Given these NAM-containing nuclear indentations, we next asked if mitochondria in confined cells were more associated to the nucleus than in suspension cells. To analyse this, we quantified NAM as mitochondria situated within the nuclear region and 10% of its surrounding area. Indeed, we found a significantly increased nucleus-mitochondria association in confined cells, which showed a consistent increase of NAM in comparison to suspension cells (Figure 2 F). The different categories of nuclear indentations contributed equally to the increase in NAM (Figure S2 C). Time-lapse imaging performed while confining down to 3 μm, showed that the confinement-induced enrichment of mitochondria at the nuclear periphery and within nuclear indentations occurred within seconds of applying mechanical stress and persisted for up to 35 minutes of confinement (Figure S2 D, Movie S1, S2). We observed no differences with respect to the amount of NAM between short or relatively longer confinement times (R^2^<0.1, Figure S2 D). This indicates that the mitochondrial redistribution within nuclear indentations is a product of the rapid and dynamic remodelling of organelle localisation which promotes a mitochondrial accumulation at the nuclear periphery.

We also confined U2OS, MDA-MB-231 and the pancreatic ductal adenocarcinoma cell line MIA PaCa-2 and quantified the amount and localization of NAM. In all cases, we found a significant increase of NAM (Figure S2 E-F), which suggests that the formation of mitochondria-containing nuclear indentation is a conserved mechanism that concentrates mitochondria at the nuclear periphery.

Overall, our results revealed that in acutely confined cells mitochondria accumulate at the nuclear periphery and within nuclear indentations.

### The ER network and actin polymerization control the accumulation of mitochondria that lead to nuclear indentations

Next, we queried how the accumulation of mitochondria around the nucleus and nuclear shape indentations are mutually controlled. Mitochondria are connected to the endoplasmic reticulum (ER)^13–15^ which is itself anchored to the nuclear membrane^16^. We hypothesised that the ER could serve as a structural scaffold that can entrap mitochondria and retain them in the perinuclear region upon confinement. Confined HeLa cells expressing a Turquoise-tagged ER-localising KDEL sequence showed an ER enrichment within the perinuclear region, which co-localised with the MitoTracker signal (Figure 3 A-C). Closer analysis, however, showed that the two signals were not completely overlapping, but rather behaved as two intertwined meshes (Figure 3 D). Line profiles of MitoTracker and KDEL-Turquoise signals spanning either the diameter of the nucleus (Figure 3 E), or shorter distances within the nuclear indentations (Figure 3 F), confirmed the presence of mitochondria and ER within nuclear indentations, organised as a network. The ER-mitochondria network dynamics regulates intracellular organelle remodelling and shape^9,17,18^. We have also previously shown that the ER network, which is a highly dynamic structure in suspension cells, gets immobilised at the nucleus-plasma membrane interface in confined cells^1^. Although the effect of cellular confinement on regulating ER immobilisation in human cells is not known, it could physically explain the anchorage of mitochondria within the nuclear region. Kymographs generated from line profiles showed a high dynamicity of ER signals in suspension cells, which was strongly reduced upon confinement (Figure 3 G-H, Movie S3), confirming our hypothesis of ER immobilisation in the nucleus-plasma membrane proximal region that can promote a nucleus-mitochondria association.

**Figure 3.**
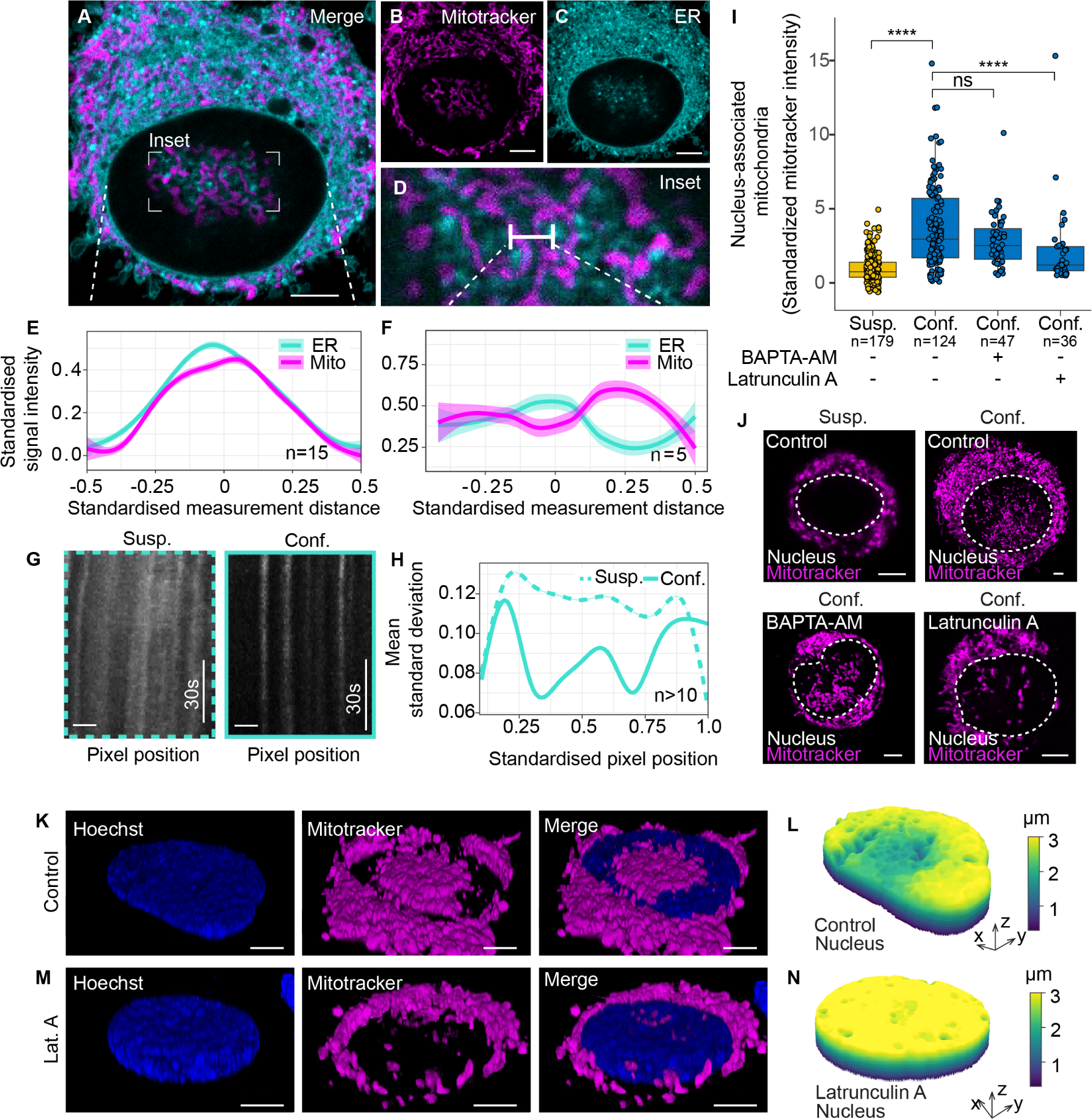
The ER network and actin polymerization control the accumulation of mitochondria at the nucleus. **(A-D)** HeLa cell expressing a Turquoise-tagged ER-localising KDEL and stained with MitoTracker. **(D)** Inset highlighted in (A). **(E)** Line profiles of ER and MitoTracker signals quantified across the nuclear diameter. Lines are mean standardised signal intensity with 95% confidence intervals. **(F)** Line profiles of ER and MitoTracker signals quantified across 1.5 μm lines starting at a point of high mitochondrial signal. Lines are mean standardised signal intensity with 95% confidence intervals. See also Figure S3 for similar analysis with late endosomes, lysosomes and peroxisomes. **(G)** Kymographs generated from whole-cell line profiles of ER signals in suspension and confined HeLa cells. **(H)** Quantification of the dynamicity of ER signal in (G) represented as the mean standard deviation with 95% confidence intervals, along the line profiles. **(I)** Quantification of NAM in HeLa cells treated with BAPTA-AM (10 μM) and Latrunculin A (500 nM) which interfere with intracellular calcium levels and actin polymerisation, respectively. **(J)** Representative MitoTracker images corresponding to BAPTA-AM and Latrunculin A treatments in (I). **(K)** 3D representation of a confined HeLa cells. Nuclei stained with Hoechst and mitochondria stained with Mitotracker. **(L)** Hoechst signal was used for an in silico reconstruction of the nucleus of the confined cell in (K). **(M)** 3D representation of a HeLa cell treated with Latrunculin A (500 nM). Nuclei stained with Hoechst and mitochondria stained with Mitotracker. **(N)** Hoechst signal was used for an in silico reconstruction of the nucleus of the Latrunculin A treated confined cell in (M). All scale bars represent 5 μm. Statistics in (I) were performed using the Wilcoxon test (* P<0.05, **** P<0.0001).

To understand whether the confinement-induced enrichment of mitochondria was specific to the mitochondria-ER network, we also stained late endosomes, lysosomes and peroxisomes in confined cells. Late endosomes showed an enrichment within the nuclear region (Figure S3 A-B), but unlike the ER, did not form a network-like structure with mitochondria (Figure S3 C). Lysosomes also exhibited similar properties as late endosomes (Figure S3 D-F). Peroxisomes appeared to be more dispersed, showing very little association to mitochondria (Figure S3 G-I).

Two major players regulating ER-mitochondrial contact sites are the actin cytoskeleton and intracellular calcium levels. Polymerization of actin filaments is required for the formation of mitochondria-ER contacts (MERCs)^19–21^, while Ca^2+^ channelling across MERCs regulates mitochondrial activity^16,20^. Therefore, using HeLa cells treated with either Latrunculin A or BAPTA-AM, we asked whether interfering with actin polymerization or intracellular calcium levels, respectively, would prevent the increase of NAM. Using immunofluorescently labelled actin and α-Tubulin in HeLa cells, we first visualised the effect of Latrunculin A or BAPTA-AM. While BAPTA-AM treatment led to a modified actin structure, Latrunculin A treatment entirely depolymerised the actin cytoskeleton and also reduced α-Tubulin structures (Figure S3 J). BAPTA-AM did not lead to a significant reduction in the overall NAM levels in the confined population, whereas Latrunculin A treatment resulted in significantly reduced levels of confinement-induced NAM (Figure 3 I-J), indicating a relevant role of actin in regulating the mitochondria-nucleus association during confinement. We further evaluated NAM accumulation in HeLa cells treated with CK666, an Arp2/3 inhibitor that impairs actin branching^22^, and SMIFH2, a Formin inhibitor that interferes with actin elongation^23^. Treatment with these inhibitors led to an altered actin cytoskeleton in terms of branching or elongation, in comparison to untreated controls (Figure S3 J). While treatment with CK666 did not lead to a reduction in NAM formation compared to control confined cells, treatment with SMIFH2 was able to reproduce the reduction in NAM observed with Latrunculin A treatment (Figure S3 K-L). Notably, treatment with Latrunculin A not only eliminated the presence of mitochondria within nuclear indentations, but also prevented the formation of nuclear shape change and nuclear indentations altogether (Figure 3 K-N). This indicates that the physical presence of the mitochondria is required for confinement-induced nuclear shape alterations and suggests a direct mechanical deformation of nuclei by the accumulated mitochondria.

Together, our data support an ER-associated and actin-mediated mechanism of NAM formation upon cellular confinement that is responsible for the formation of nuclear indentations.

### NAM fuel a nuclear ATP surge upon confinement

Functional analysis of proteins enriched in the nuclear fraction after confinement (Figure 1 B) revealed biological processes that regulate ATP synthesis (Figure 4 A). We hypothesised that mitochondria within nuclear indentations may actively produce energy to help cells and the nucleus compartment to adapt to mechanical confinement stress. To test if confinement-induced NAM formation could influence nuclear ATP levels, we performed Förster resonance energy transfer (FRET) imaging using cells transiently expressing a nuclear ATP sensor^24^. We found a significant and rapid increase in nuclear ATP levels in HeLa cells upon confinement (Figure 4 B-C). Inhibition of mitochondrial ATP synthesis using Oligomycin A prevented this nuclear ATP surge observed in confined HeLa cells (Figure 4 B-C) but did not alter the actin cytoskeleton (Figure S4 A), indicating that mitochondrial ATP production is responsible for this nuclear ATP increase. Treatment of HeLa cells with Latrunculin A, which interfered with the confinement-induced increase in NAM, also abolished the increase in nuclear ATP in confined cells, confirming NAM as the main ATP source (Figure 4 B-C). As previously discussed with regards to the dynamics of NAM formation (Figure S2 D), the duration of mechanical cell confinement also did not influence the nuclear ATP increase, which was an immediate response and persisted for at least up to 35 minutes of confinement (Figure S4 B).

**Figure 4.**
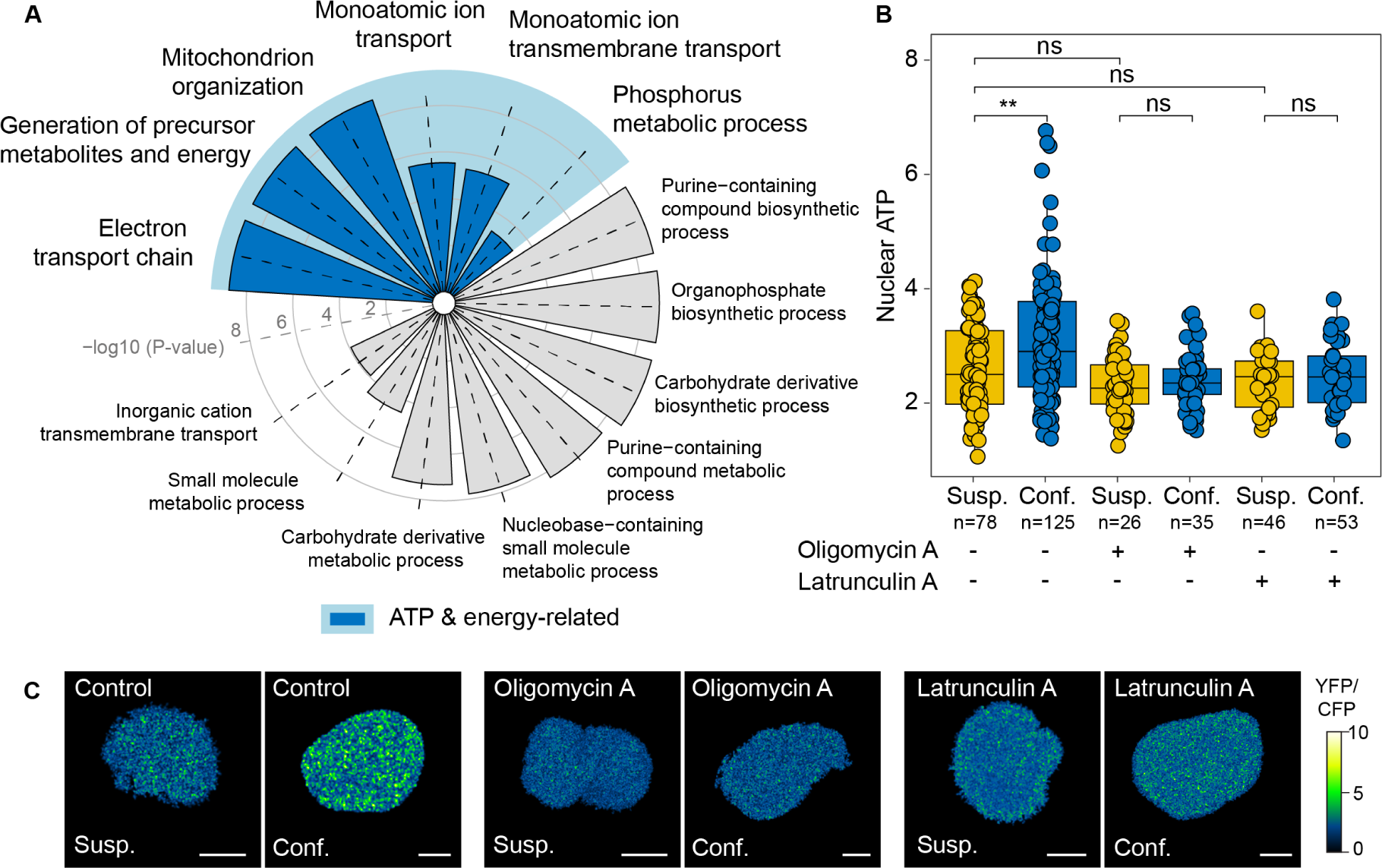
Confinement induces a NAM-fuelled nuclear ATP surge. **(A)** Biological processes – GSEA of proteins enriched in the confined nuclear fraction. Concentric circles represent -log10 (P-value). **(B)** Nuclear ATP quantification in HeLa cells treated with Oligomycin A (1 μM) or Latrunculin A (500 nM) using a FRET-based nuclear ATP sensor^24^. **(C)** Representative images of nuclear ATP in HeLa cells showing corresponding FRET ratios as quantified in (B). All scale bars represent 5 μm. Statistics in (B) were performed using the Wilcoxon test (* P<0.05, ** P<0.01). See also Figure S4 for related experiments in U2OS cells.

To verify if confinement induced a nuclear ATP surge in a cell line-independent manner, we next used U2OS cells and measured confinement-dependent nuclear ATP levels. Similar to our observations in HeLa cells, mechanical confinement triggered a nuclear ATP increase in U2OS cells, which could be blocked by inhibiting mitochondrial ATP synthesis through Oligomycin A treatment (Figure S4 C-D). Finally, we queried whether this increase in nuclear ATP was attributed to increased mitochondrial ATP production upon confinement. Using a FRET-based mitochondrial ATP sensor^24^ transfected into HeLa cells, we did not observe any confinement-dependent differences in mitochondrial ATP production either when analysing all mitochondria or only NAM (Figure S4 E-G).

Overall, these results suggest that even though confinement does not trigger general mitochondrial ATP production, the tight physical contact between the NAM and the nucleus at nuclear indentations, is responsible for a nuclear ATP surge in confined cells.

### Nuclear ATP regulates chromatin compaction under confinement

ATP plays an important role as a biological hydrotrope^25^. Nuclear ATP has previously been hypothesised to regulate chromatin condensation by influencing free Mg^2+^ ion levels^26^. Upon hydrolysis of ATP, Mg^2+^ ions are transiently released leading to chromatin condensation, for instance, during cell division^27^. Conversely, high ATP levels promote ATP-Mg^2+^ complexes, which may result in increased chromatin accessibility and solubility^26^. In the context of mechanical stress, this increased accessibility, mediated by the loss of heterochromatin, results in nuclear softening, thereby insulating genetic material against mechanical force^28^.

To evaluate the chromatin state between suspension and acutely confined HeLa cells, we used Hoechst signal to quantify the nuclear coefficient of variation (Figure 5 A-B). We observed a nominally higher coefficient of variation upon confinement (Figure 5 B). Treatment with Oligomycin A did not alter the coefficient of variation in suspension cells but showed a higher coefficient of variation in confined cells (Figure 5 A-B). To evaluate how the coefficient of variation was associated to the spatial organization of condensed chromatin structures, we performed an iterative binning and calculated the coefficient of variation for increased pixel-based binning areas. Binning image pixels reduces smaller and local signal variations and enables efficient computational comparisons across larger and more distinct chromatin structures. While control suspension, control confined and Oligomycin A treated suspension cells showed a sharp decline in their respective coefficient of variation for increased binning sizes, Oligomycin A treated confined cells continued to show a high coefficient of variation (Figure 5 A-B), suggesting that the absence of ATP upon confinement maintains spatial heterogeneities in chromatin compaction and prevents chromatin solubility.

**Figure 5.**
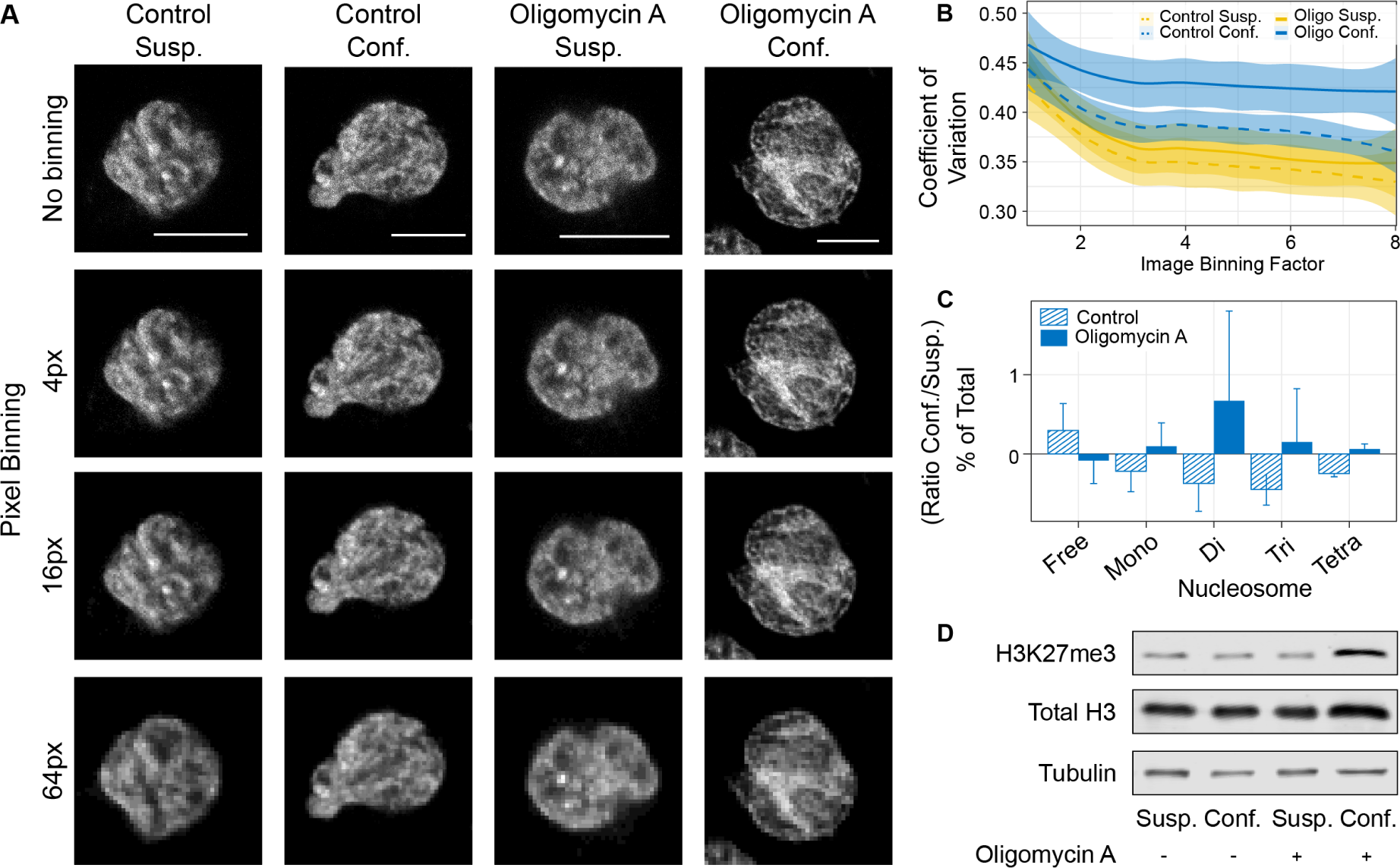
Nuclear ATP regulates chromatin compaction. **(A)** Hoechst staining in HeLa cells in suspension or confined, and either untreated or treated with Oligomycin A (1 µM). Signal intensities were binned progressively starting from no binning, and up to 8x8 (64 pixels) bins. **(B)** Quantification of coefficient of variation (standard deviation / mean) of Hoechst signal from nuclei as in A. Binning was done for X^2^ pixels, X = {1 … 8; increment = 1} **(C)** Free, mono-, di-, tri- or tetra-nucleosomes, as a percentage of the total nucleosome count in control or Oligomycin A (1 µM) treated HeLa cells. Data is represented as the ratio between confined and suspension cells. **(D)** Western blot of H3K27me3 facultative heterochromatin mark, total histone H3, and loading control α-Tubulin in Control or Oligomycin A (1 µM) treated, suspension or confined HeLa cells. All scale bars are 10 µm.

To further assess chromatin accessibility and condensation, we performed an assay for transposase-accessible chromatin with sequencing (ATAC-Seq) on control and Oligomycin A treated suspension and confined cells. A principal component analysis validated the clustering of replicates but showed little separation between suspension and confined conditions (Figure S5 A). The evaluation of differential accessibility between suspension and confined conditions, in both control and Oligomycin A treated samples led to negligible functional changes (Figure S5 B). We hypothesized that following acute confinement, chromatin does not initially reorganize to allow for specific transcriptional programs, but rather as a physical adaptation to absorb the mechanical force. This would lead to stochastic chromatin relaxation rather than changes in the accessibility of gene regulatory regions. To test this hypothesis, we quantified the percentage of nucleosome-free, mono-nucleosome, di-nucleosome, tri-nucleosome and tetra-nucleosome regions (Figure S5 C). Confinement, in control conditions, induced an increase in nucleosome-free regions and a decrease in nucleosome-containing regions compared to suspension cells (Figure 5 C, Figure S5 C), suggesting increased chromatin relaxation and accessibility. In contrast, compared to suspension cells, Oligomycin A treated cells displayed more nucleosome-containing regions and less nucleosome-free regions during confinement (Figure 5 C, Figure S5 C). This suggests that nuclear ATP plays a crucial role in regulating chromatin relaxation as a response to mechanical stress.

Chromatin accessibility and organisation is typically regulated by epigenetic modifications such as tri-methylation on histone H3 lysine residues, which are regarded as chromatin repressive marks, reducing accessibility and promoting condensation^29^. Indeed, Oligomycin A treated cells which showed low levels of chromatin relaxation, exhibited remarkably high levels of H3K27me3, a facultative heterochromatin mark (Figure 5 D). Taken together, our data shows the importance of a nuclear ATP surge upon confinement which is vital for chromatin relaxation, and hence nuclear softening^28^, in response to mechanical stress.

### Nuclear ATP is required for timely DNA damage repair and S-phase progression

Confinement can induce DNA damage^30^, which, if not repaired, can impair DNA replication and cell cycle progression^31,32^. Moreover, increased chromatin accessibility is an important pre-requisite for efficient DNA damage repair^33^, as is high amounts of ATP^34–36^, together providing access to DNA repair machinery and the required energetics. Hence, we hypothesised that in addition to regulating chromatin solubility in response to mechanical stress, the nuclear ATP surge observed during confinement may be necessary to ensure that cells have sufficient ATP to promptly repair damaged DNA following confinement.

Using U2OS cells expressing a truncated-53BP1 reporter^37^, we confirmed that confinement induces an immediate increase of DNA damage foci, which were absent in cells treated with Oligomycin A (Figure S6 A-B). DNA damage repair signalling, through 53BP1 recruitment and respective downstream events are triggered by phosphorylation and therefore require ATP^34–36^. As a result, we hypothesised that the lack of 53BP1 foci detected in confined Oligomycin A treated cells may be due to the inability of cells to activate DNA damage repair signalling, rather than the prevention of DNA damage itself. To test this hypothesis, we released cells from confinement and Oligomycin A treatment, and quantified the accumulation of 53BP1 foci over time following cell adhesion (Figure 6 A). 3 hours after confinement release, U2OS cells which had been subjected to acute confinement, were fully adhered and showed trends of higher levels of 53BP1 foci (Figure S6 C). These foci returned to baseline levels 1 hour later, in agreement with the reported dynamics of 53BP1^38^. Contrastingly, and in accordance with our hypothesis, while no DNA damage was observed in Oligomycin A treated cells during confinement, 3 hours after confinement and Oligomycin A release, cells showed significantly increased levels of 53BP1 foci. These foci persisted for up to 7 hours, exhibiting a 3-hour delay in DNA damage repair compared to untreated confined cells (Figure S6 C). These data confirm that the lack of a confinement-induced nuclear ATP surge prevents timely DNA damage signalling and repair.

**Figure 6.**
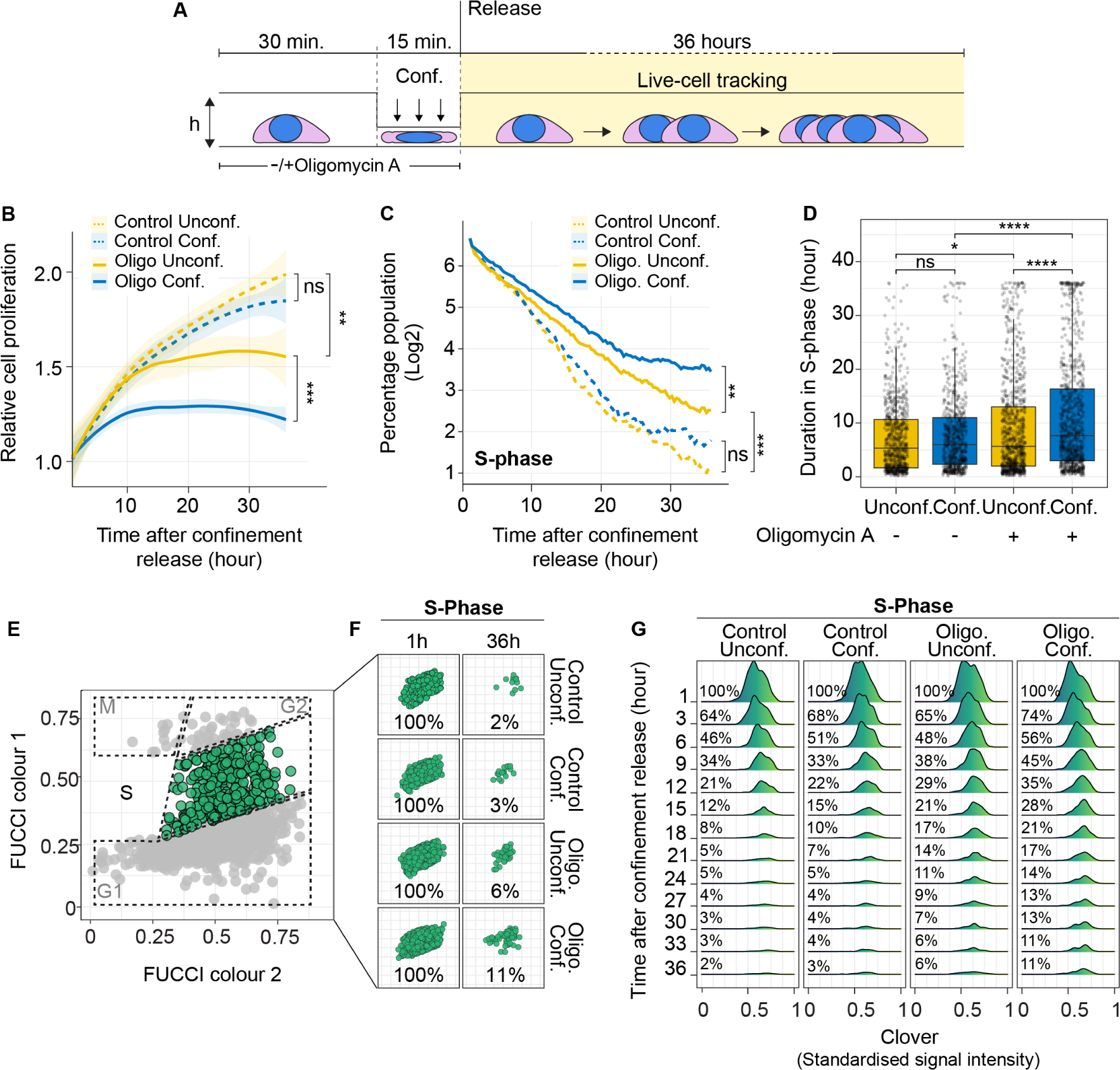
Nuclear ATP surge during confinement is required to safeguard S-phase progression after confinement release. **(A)** Schematic representation of the protocol used to study the effect of acute confinement on cells after release. Briefly, U2OS-FUCCI cells are given a 30-minute treatment of Oligomycin A, following which they are acutely confined for 15 minutes. Confinement and Oligomycin A treatment are then released, cells allowed to attach under normal growth conditions and tracked (as single cells) for 36 hours to study cell proliferation or FUCCI-based cell cycle progression. **(B)** Relative cell proliferation of U2OS cells that were either unconfined or confined, and either left untreated or treated with Oligomycin A (1 μM). **(C)** Time taken by cells detected in the S-phase after confinement release, indicating the beginning of tracking, to progress through the S-phase. **(D)** Progression of cells detected in the S-phase after confinement release, at the beginning of tracking, for 36 hours. **(E)** Single cell analysis for FUCCI is performed by spatially distributing cells based on standardised signal intensity (see methods). Population highlighted in green represents the S-phase. **(F)** Single cell representation of cells first detected in the S-phase after confinement release at the first (1^st^ hour) and last (36^th^ hour) time points. **(G)** Progression of the S-population at 3-hour intervals. Statistics in (B) and (D) was performed using the Mann-Whitney-Wilcoxon test (* P<0.05, ** P<0.01, *** P<0.001). Statistics in (C) was performed using the Wilcoxon test (* P<0.05, **** P<0.0001). See also Figure S5 for details on the FUCCI system and, G1-phase and G2-phase dynamics.

The progression of the cell cycle, which is the basis for cell proliferation, is tightly regulated by cellular energy homeostasis^26^, and dependent on efficient DNA damage repair, especially during the S-phase of the cell cycle^29–31^. During the cell cycle, ATP synthesis is boosted at G1-S-phase transition, when mitochondria fuse into long branches of hyperpolarized ATP factories^27^. This increase of ATP is required to activate Cyclin-E and the progression into S-phase, which is estimated to require 4-5 mM of nuclear ATP levels for optimal speed^27^.

Hence, we questioned whether the observed delay in DNA damage repair would affect proliferation in cells that had been confined in the presence of Oligomycin A. This would indicate that mechanical stress negatively affects cell fitness in the absence of the confinement-induced nuclear ATP surge. To assess this, U2OS cells were acutely confined for 15 minutes, with or without a short 30-minute Oligomycin A treatment, and then returned to standard culture conditions where they were tracked for 36 hours (Figure 6 A). Exposure to this brief mechanical confinement by itself did not significantly alter proliferation of U2OS cells when compared to unconfined control cells over 36 hours (Figure 6 B). This suggests that acute confinement does not result in mechanical stress-induced memory that impairs cell proliferation of future generations. Contrastingly, a short 30-minute Oligomycin A treatment alone significantly reduced cell proliferation, arguing for the perpetration of a metabolic-stress memory that impacts fitness. Notably, this short 30-minute Oligomycin A treatment when followed by an acute 15-minute confinement, almost entirely abolished cell proliferation (Figure 6 B). This suggests that acute metabolic stress sensitises cells towards acquiring a strong mechanical stress-induced memory that arrests cell proliferation.

To further explore the effect of acute confinement on cell proliferation over time after confinement release, we used an adapted FUCCI system (Figure S6 D)^39,40^ which facilitates live cell tracking through all the cell cycle phases (Figure S6 E). For each condition, we divided the cell population into the different phases – G1, S and G2 – in which they were first detected upon confinement release, when we started the tracking. We analysed the progression of these 100% pure G1, S or G2 populations, from the beginning of tracking, into their respective next cell cycle phases. This allowed us to identify phase-specific effects of either confinement, Oligomycin A treatment, or their combination. We observed that while confinement or Oligomycin A treatment alone had a significant impact on the progression of G1 (Figure S6 F) and G2 (Figure S6 G) phases, there was little to no additional effect when combining the two treatments. These results suggest that confinement-induced nuclear ATP surge is not critical for G1 and G2 phase progression.

Given the observed delay in DNA damage repair, we queried whether the S-phase of the cell cycle could be particularly affected by the combination of a short Oligomycin A treatment and acute confinement. Upon analysis of cells that were in the S-phase at the beginning of tracking, we observed that acute confinement alone did not delay the progression of cells into the next phase (Figure 6 C). Oligomycin A treatment, on the other hand, resulted in a significant delay in S-phase progression, which was exacerbated in conjunction with acute confinement (Figure 6 C). Quantification of S-phase duration corroborated these findings (Figure 6 D). This S-phase-specific impact of Oligomycin A treatment together with acute confinement may explain the observed defect in cell proliferation (Figure 6 B). As a consequence, we observed a larger population of cells remaining in the S-phase at the end of the 36-hour tracking in Oligomycin A treated cells, which was further increased upon the addition of acute confinement (Figure 6 E-F). This additive effect of short Oligomycin A treatment and acute confinement on S-phase progression was especially evident from the beginning of tracking, resulting in 10% more cells retained in the S-phase (Figure 6 G). By the end of the 36-hour tracking, control unconfined cells showed only 2-3% of the initial S-phase population remaining in the S-phase, with acute confinement having almost no effect on this remaining population (Figure 6 G). Contrastingly, while Oligomycin A treatment increased this remaining population by approximately 2-fold, the combination of a short Oligomycin A treatment and acute confinement together led to a 4-fold increase in the remaining S-phase population (Figure 6 F-G).

These data indicate that both confinement and Oligomycin A treatment influence cell cycle progression by altering the duration of the G1 and the G2 phases. However, the combination of the two treatments, in which the nuclear ATP surge is blocked during confinement, notably affects S-phase progression, likely as a consequence of impaired DNA damage repair.

## DISCUSSION

Recent work has highlighted the importance of the nucleus as a hub for mechanotransduction, regulating adaptation dynamics to mechanical stress^11,28,41,42^. However, the effect of mechanical stress in the form of confinement on the subcellular distribution of proteins and metabolites remains entirely unknown.

Intracellular mitochondrial re-localisation under stress conditions enables energetically intensive processes such as cancer cell migration through crowded tumour microenvironments^43,44^, and cellular adaptation through mitochondria-nucleus contact sites^45^. In this study, we coupled nucleo-cytoplasmic subcellular fractionation with mass spectrometry, to identify an enrichment of mitochondrial proteins in the nuclear compartment during confinement. High-resolution microscopy confirmed the accumulation of mitochondria at the nuclear periphery (NAM), resulting in nuclear shape change in the form of indentations. This NAM formation is actin-dependent and co-localises with an immobilised ER network. The ER and mitochondria have been shown to co-regulate stress responses where actin-mediated MERCs are critical to maintain cellular homeostasis^20,21^. Our data further pointed towards altered biological processes such as electron transport chain and energy production, leading to our discovery of a nuclear ATP surge upon mechanical stress in confinement. In germ cell development, recent work has highlighted the ability of mechanical stimuli from surrounding tissue to activate mitochondrial metabolism^46^. We were also able to interfere with the mechanical stress induced increase of nuclear ATP levels by independently inhibiting mitochondrial ATP synthesis, or actin depolymerisation, emphasising the relevance of nucleus-mitochondria proximity. Indeed, actin, a key regulator of cellular mechanics, plays a pivotal role in responding to mechanical cues for mitochondrial dynamics together with the ER, and has been shown to regulate mitochondrial fission^18^, cellular homeostasis through mitochondrial turnover and mitophagy^14^, cytokinesis during cell division^19^, and mitochondrial transport through association with mitochondrial myosins^47^.

Among the many instances of mechanical confinement-induced stress in development and diseases^48–50^, cancer cells must overcome important mechanically confining processes that challenge cell fitness and survival, calling for buffering mechanisms against mechanical stress. For instance, compressive mechanical stress has been shown to regulate cancer cell spheroid growth, by inducing a G1 arrest due to volume limitation^51^, or causing cell proliferation arrest through apoptosis^52^. Previous work has also highlighted that nuclear ATP levels regulate DNA duplication^53,54^, chromatin re-organisation and DNA damage repair^26^, all of which are critical processes for S-phase progression.

Our study, notably, highlights the short-and long-term response of cancer cells to acute mechanical stress. Confinement, within a matter of seconds, induces a rapid NAM formation and the consequent nuclear shape changes, and increase in nuclear ATP which is required for chromatin solubility^26^ to insulate against DNA damage from mechanical forces^28^. Pharmacological inhibition of the nuclear ATP surge significantly reduced chromatin accessibility, evidenced by a high number of nucleosomes and heterochromatin post-translational modifications. In fact, without proper accessibility, DNA damage repair is inefficient, and hence chromatin epigenetic marks that are associated with active transcription are also often shared with repair processes^55^. Consequently, we also observed a marked delay in the repair of confinement-induced DNA damage in cells that were subject to the pharmacological inhibition of nuclear ATP upon confinement, which in the long-term, impacts post-confinement cell proliferation, with specific consequences for cells confined in the S-phase. In essence, acute metabolic stress sensitises cells towards acquiring a mechanical stress-induced memory that leads to proliferation defects. Our results highlight the metabolic adaptive response of cancer cells to acute stress, and how just a 15-minute confinement is sufficient to detriment post-confinement fitness when the nuclear ATP increase is blocked.

Overall, our work highlights the importance of a physical association of mitochondria to the nucleus as an ATP source, which acts as a safeguard for proper cell cycle progression.

## ACKNOWLEDGMENTS

We would like to thank the CRG Advanced Light Microscopy Unit for their support with imaging experiments and the ICFO Nanofabrication Laboratory for the design and production of moulds necessary for generating confinement coverslips. We also thank Anamaria Elek and Arnau Sebé-Pedrós for their insightful discussions and exploratory analysis. Last, but not least we thank all the Sdelci and Ruprecht lab members for their discussions and critical feedback on this manuscript.

## AUTHOR CONTRIBUTIONS

Conceptualization: RG, SS, VR

Methodology: RG, FP

Software: RG, SK, IS, AGZ, UG

Validation: RG, FP

Formal analysis: RG, FP, SK, IS, SD, AGZ, UG, VV, KP, ACM

Resources: SK, IS, KP, ACM

Investigation: RG, FP, AGZ, LE, KP

Data curation: RG, FP, SK, KP, ACP

Writing – original draft: RG, SS, VR

Writing – review & editing: RG, FP, AGZ, UG, SS, VR

Visualisation: RG, FP

Supervision: SS, VR

Project administration: RG, SS, VR

Funding acquisition: SS, VR

## FIGURES & FIGURE LEGENDS

**Figure S1.**
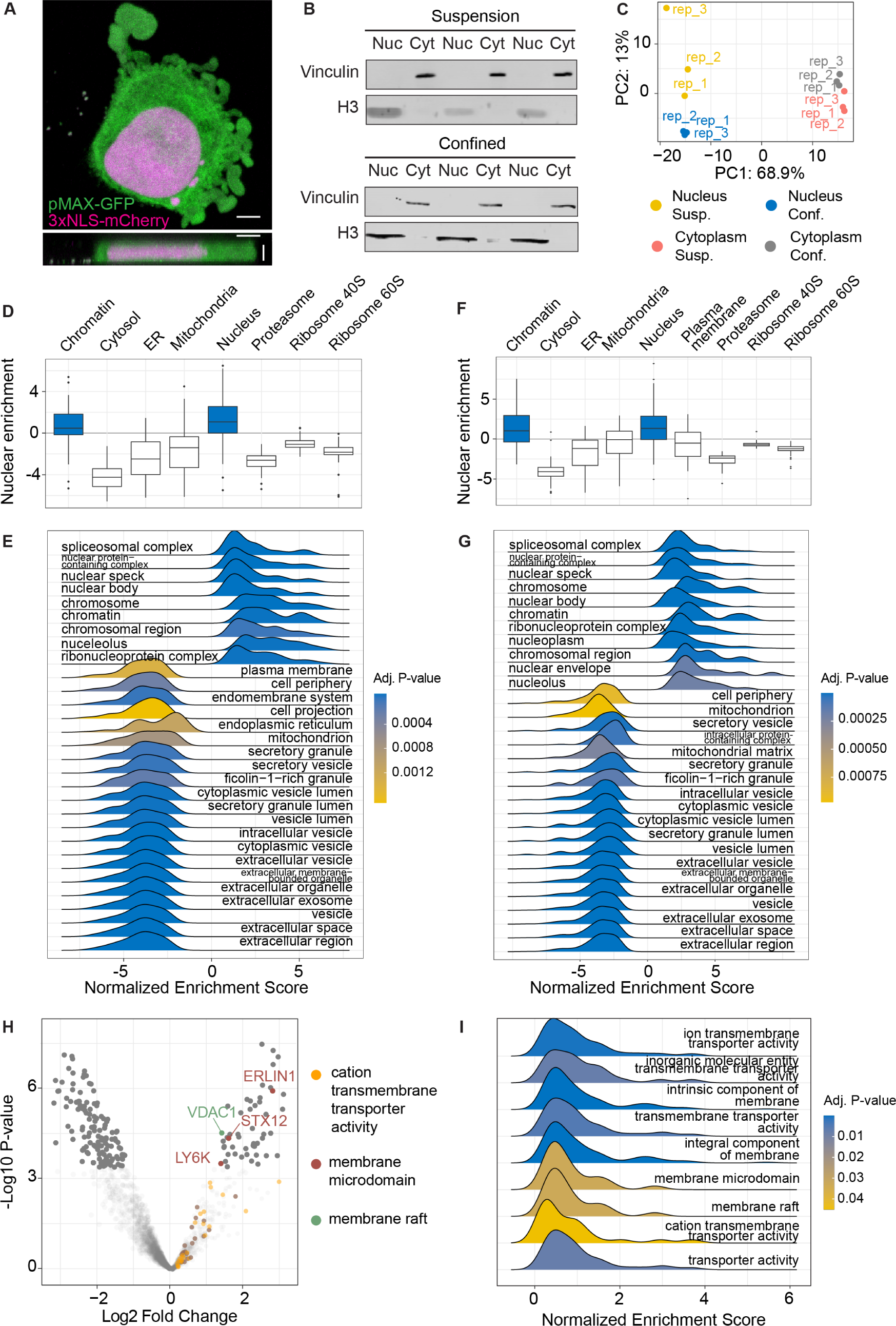
Related to Figure 1. Quality control for subcellular fractionation and proteomics. **(A)** HeLa cells displaying blebs when confined to a height of approximately 3 μm using the agarose confiner. **(B)** Western Blot to validate nucleo-cytoplasmic fractions of samples obtained through subcellular fractionation of HeLa cells in suspension or under confinement. H3 antibody and Vinculin antibodies were used to test nuclear or cytoplasmic fractions, respectively. **(C)** PCA of results obtained from MS showing clustering among replicates and a clear separation between and within fractions (nucleus vs cytoplasm) and conditions (confinement vs suspension). **(D)** Enriched and depleted terms when comparing the nuclear fraction of suspension HeLa cells to its cytoplasmic fraction. Enrichment of chromatin and nuclear terms, and depletion of cytosolic and other organelle terms confirms successful nucleo-cytoplasmic fractionation of suspension cells. **(E)** GSEA of proteins from the nucleus-cytoplasm comparison in (D). **(F)** Enriched and depleted terms when comparing the nuclear fraction of confined HeLa cells to its cytoplasmic fraction. Enrichment of chromatin and nuclear terms, and depletion of cytosolic terms confirms successful nucleo-cytoplasmic fractionation of cells under confinement. **(G)** GSEA of proteins from the nucleo-cytoplasmic comparison in (F). **(H)** Enriched and depleted terms when comparing the cytoplasmic fraction of confined HeLa cells to that of unconfined cells. **(I)** Comparison of GSEA performed on the cytoplasmic fractions in (H) only revealed membrane transport activity-related terms. All horizontal scale bars represent 5 μm. Vertical scale bar in A represents 3 µm.

**Figure S2.**
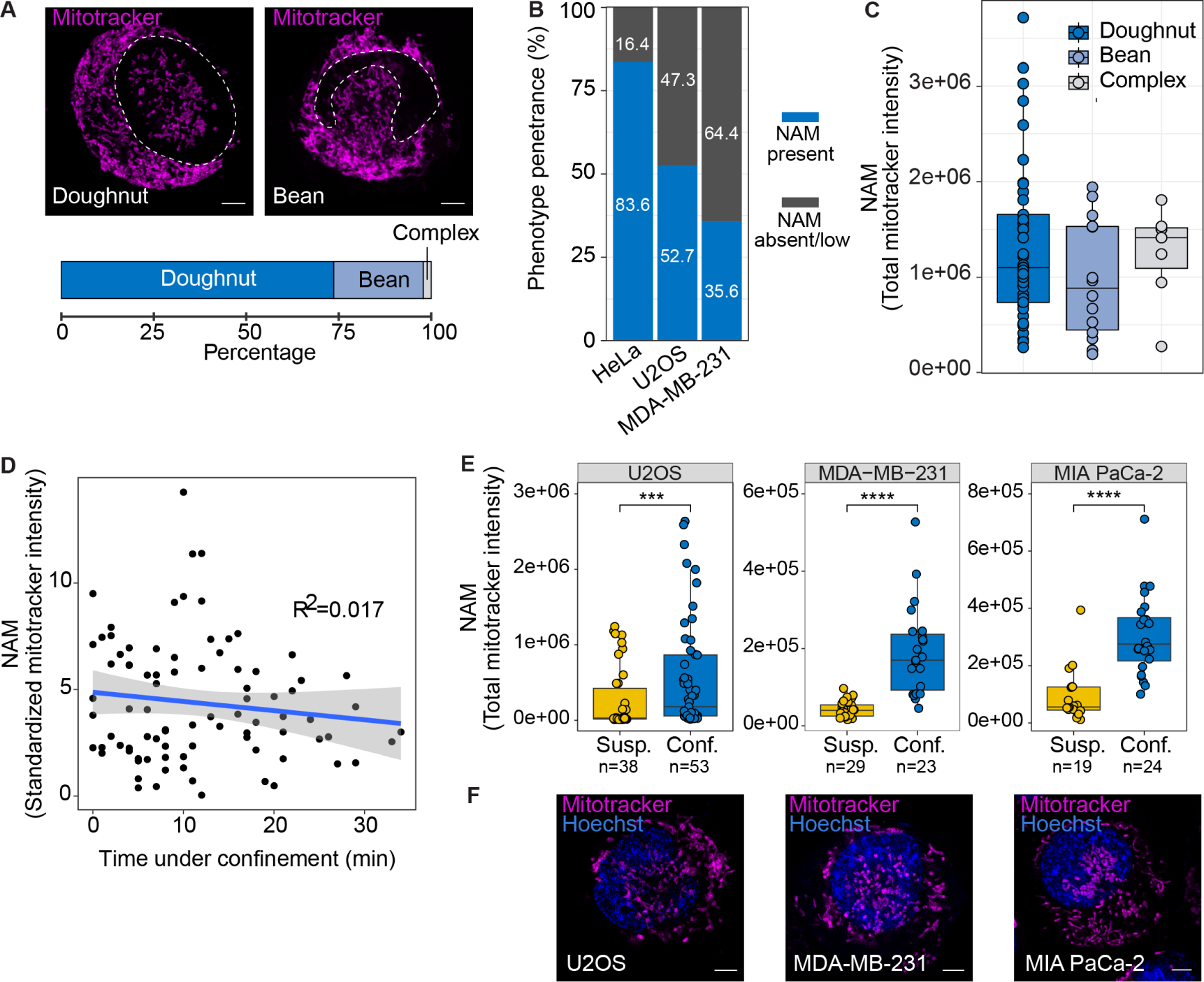
Related to Figure 2. Characterisation of NAM and nuclear indentations. **(A)** [top] Nuclear indentations were classified into either “doughnuts”, “beans”, or complex nuclei based on the Hoechst signal and localisation of mitochondria. [bottom] Quantification of nuclear indentations as a percentage of the population positive for nucleus-associated mitochondria (NAM). **(B)** Penetrance of the NAM phenotype in different cell lines. **(C)** Correlation of NAM levels with nuclear shape classification in (A). **(D)** Correlation of NAM levels in HeLa cells with the duration of confinement. **(E)** Quantification of NAM in osteosarcoma U2OS cells, triple-negative breast cancer MDA-MB-231 cells, and pancreatic ductal adenocarcinoma MIA PaCa-2 cells. **(F)** Representative images of confined cell lines quantified in (E), stained with Hoechst (nucleus) and MitoTracker (mitochondria). All scale bars represent 5 μm. Statistics in (E) were performed using the Wilcoxon test (* P<0.05, *** P<0.001, **** P<0.0001).

**Figure S3.**
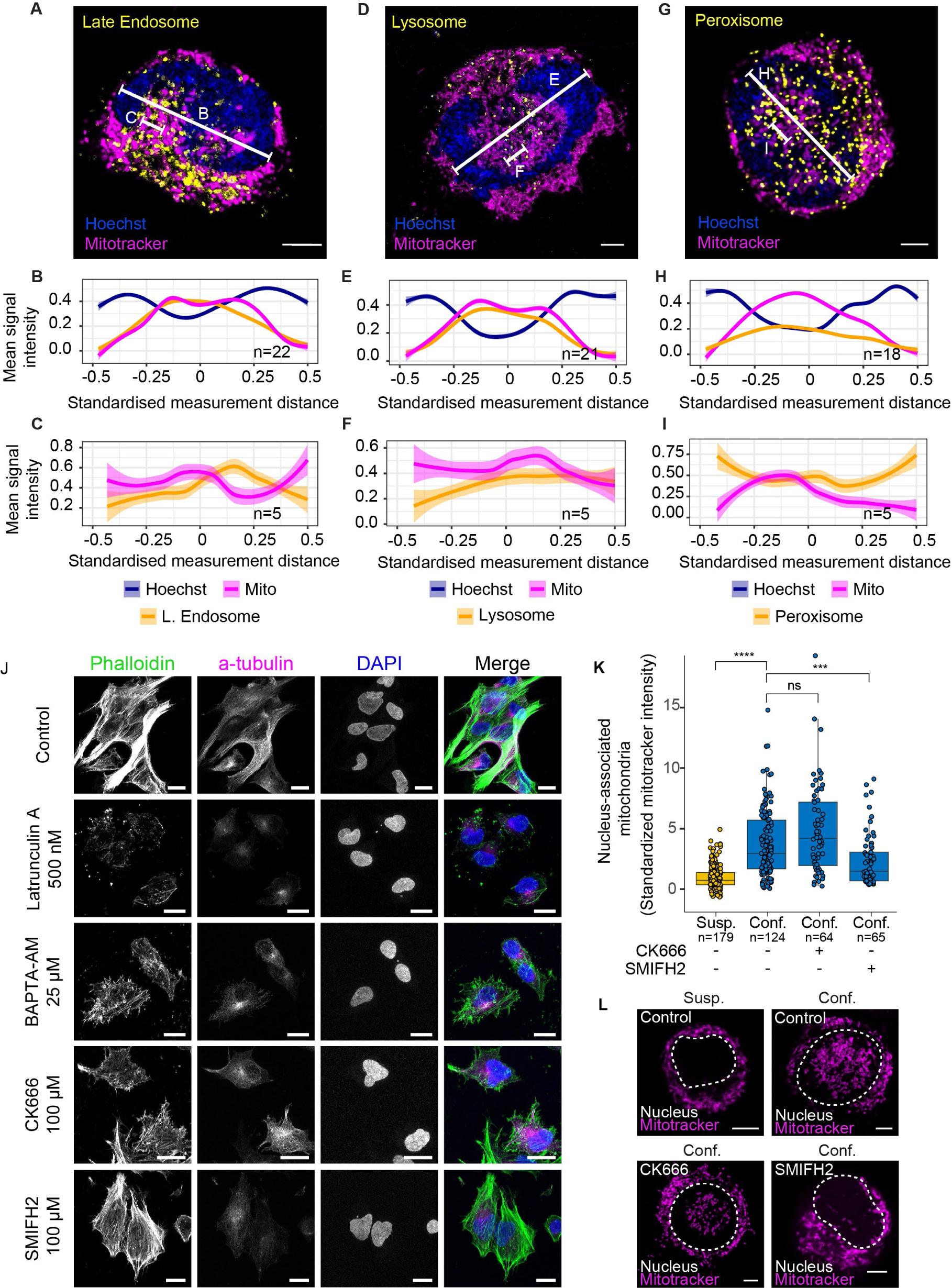
Related to Figure 3. Mitochondrial co-localization with various organelles. **(A-I)** Comparison of mitochondrial signal with the localization of late endosomes (A-C), lysosomes (D-F) and peroxisomes (G-I). **(B, E, H)** Line profiles of MitoTracker and respective organelles quantified across the nuclear diameter as indicated in their representative images. **(C, F, I)** Line profiles of ER and MitoTracker signals quantified across 1.5 μm lines starting at a point of high mitochondrial signal. **(J)** Immunofluorescent images of fixed adherent HeLa cells stained with Phalloidin (marks actin), α-Tubulin, or DAPI. Cells were treated with Latrunculin A (500 nM), BAPTA-AM (25 µM) or CK666 (100 µM) for 30 min, or SMIFH2 (100 µM) for 1 hour. **(K)** Quantification of NAM in HeLa cells treated with CK666 (100 μM) and SMIFH2 (100 µM) which interfere with actin branching and actin polymerisation/elongation, respectively. **(L)** Representative MitoTracker images corresponding to CK666 and SMIFH2 treatments in (K). Scale bars in (A), (D), (G) and (L) represent 5 μm. Scale bars in (J) represent 20 µm. Statistics in (K) were performed using the Wilcoxon test (* P<0.05, *** P<0.001, **** P<0.0001).

**Figure S4.**
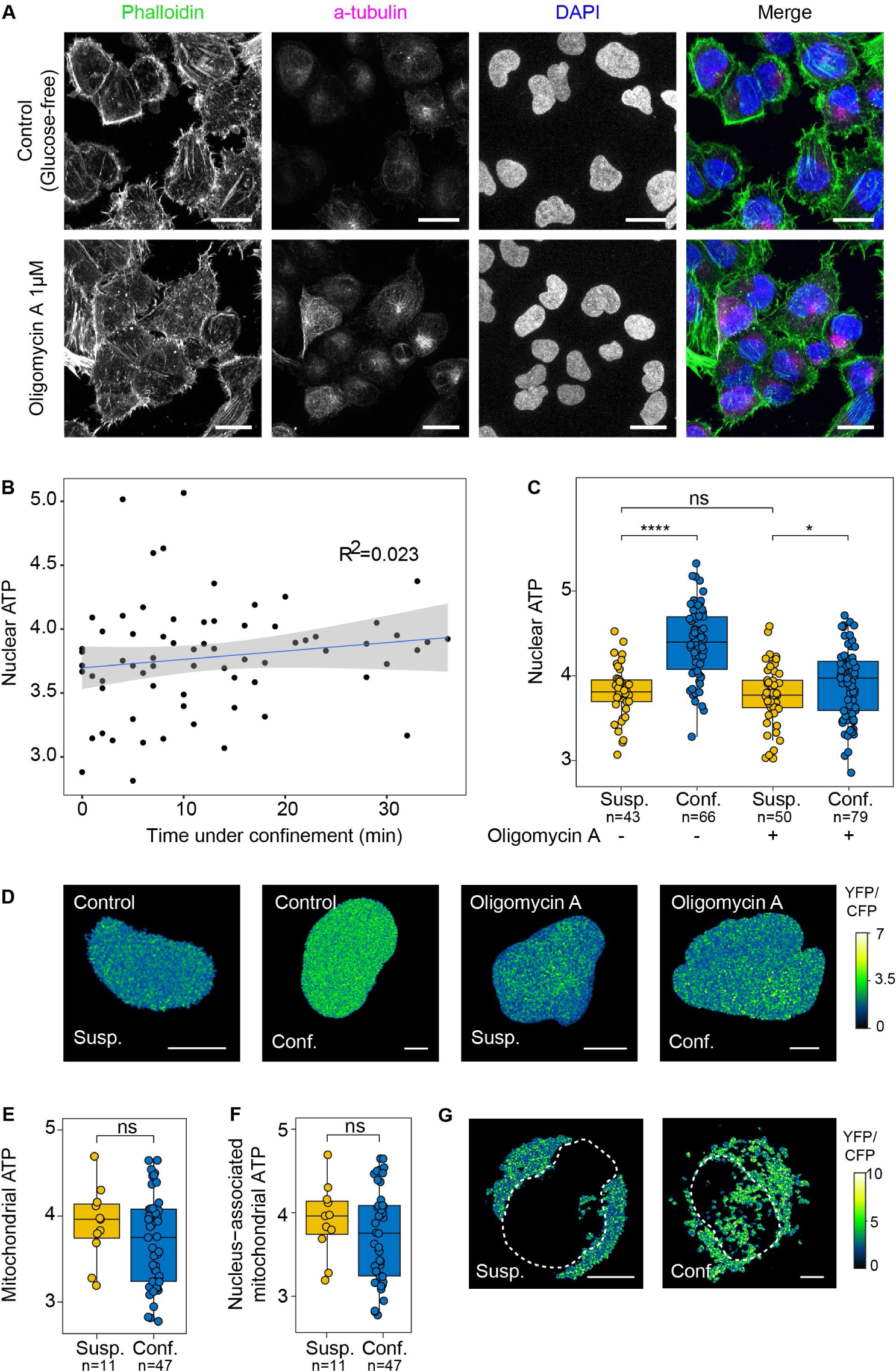
Related to Figure 4. Nuclear and mitochondrial ATP. **(A)** Immunofluorescent images of fixed adherent HeLa cells stained with Phalloidin (marks actin), α-Tubulin, or DAPI. Cells were treated with Oligomycin A (1 µM) for 30 min. **(B)** Correlation of nuclear ATP levels in HeLa cells with the duration of confinement. **(C)** Nuclear ATP quantification in U2OS cells treated with Oligomycin A (1 μM) using a FRET-based nuclear ATP sensor. **(D)** Representative images of nuclear ATP in U2OS cells showing corresponding FRET ratios as quantified in (C). **(E-F)** Quantification of whole-cell mitochondrial ATP levels (E) and NAM-specific ATP levels (F) in suspension and confined HeLa cells using a FRET-based mitochondrial ATP sensor. **(G)** Representative images of mitochondrial ATP in HeLa cells showing corresponding FRET ratios as quantified in (E-F). Scale bars in (A) represent 20 µm. All other scale bars represent 5 μm. Statistics in (C), (E) and (F) were performed using the Wilcoxon test (* P<0.05, **** P<0.0001).

**Figure S5.**
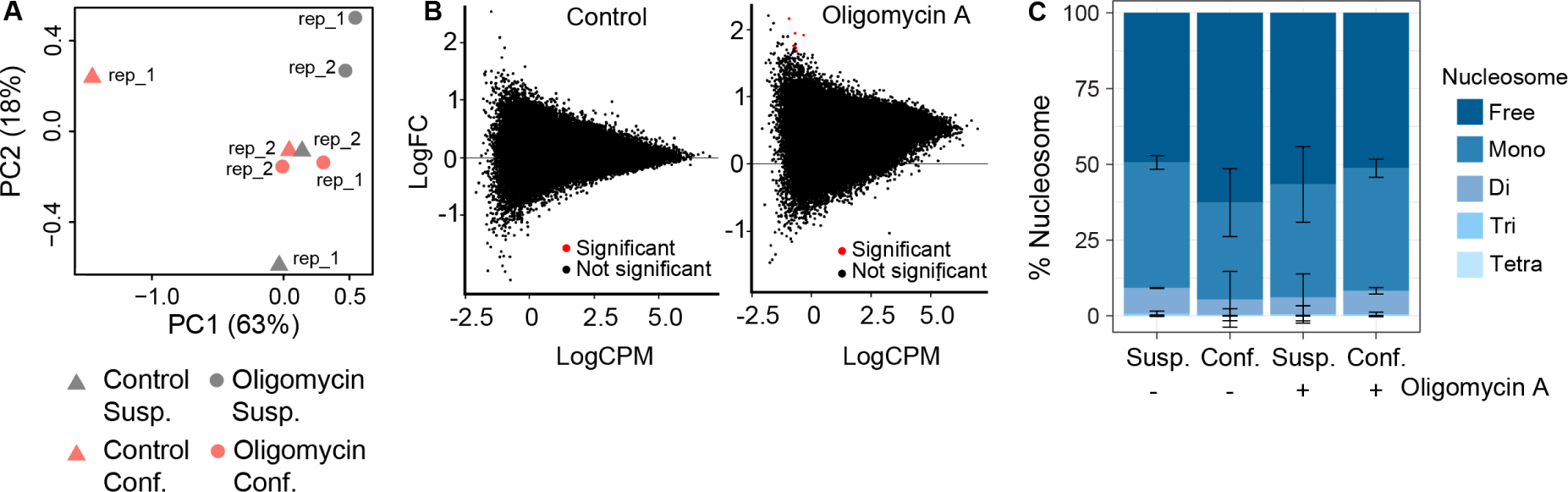
Related to Figure 5. Nuclear ATP regulates chromatin compaction under confinement. **(A)** Principal component analysis of ATAC sequencing data for suspension or confined HeLa cells either treated or untreated with Oligomycin A (1 µM). **(B)** M-A plot of ATAC sequencing data for HeLa cells untreated or treated with Oligomycin A (1 µM). Data is shown as confinement vs. suspension, in either control untreated or Oligomycin A treated cells. **(C)** Free, mono-, di-, tri-or tetra-nucleosome count as a percentage of the total nucleosome count from ATAC sequencing data as in (A-B).

**Figure S6.**
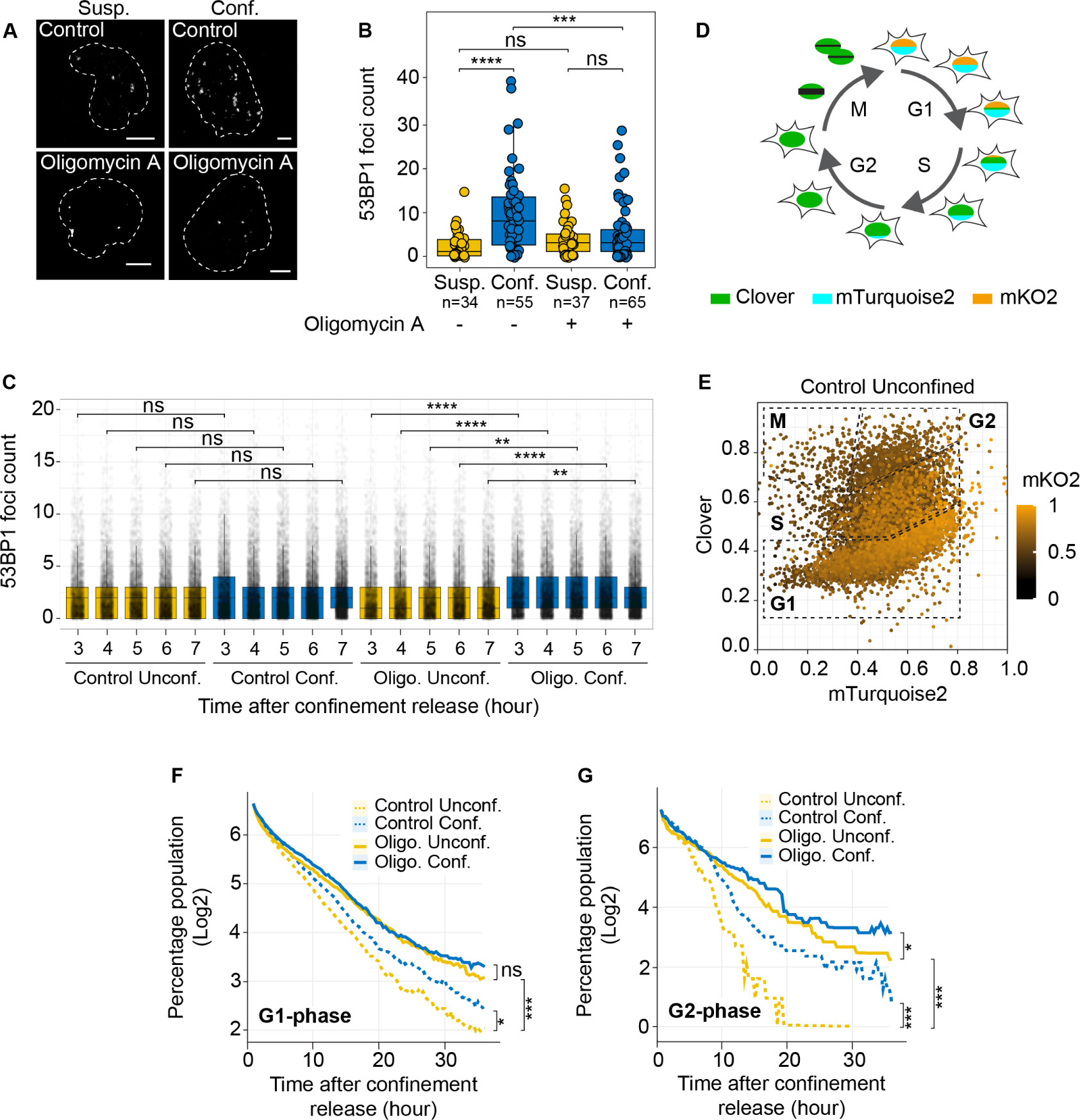
Related to Figure 6. DNA damage and cell cycle. **(A)** Visualisation of the DNA damage repair enzyme 53BP1 using U2OS cells expressing a truncated-53BP1-maroon reporter. Images are maximum intensity projections, and the outline of the nucleus is based on transmitted light images. **(B)** Quantification of 53BP1 foci in cells in suspension or under confinement, untreated or treated with Oligomycin A (1 µM). Quantification is performed on maximum intensity projection images as in (A). **(C)** Quantification of 53BP1 foci in U2OS cells post-confinement release. **(D)** FUCCI cells express clover, turquoise and orange (mKO2) fluorophores tagging cell cycle proteins (SLBP-Turquoise2, Clover-Geminin, Cdt1-mKO2). **(E)** Specific combinations of expression levels of these fluorophores indicate the cell cycle phase. For analysis during this study, gating for phases was done using an XYZ projection (x axis = SLBP-Turquoise2, y axis = Clover-Geminin, z axis plotted as colour = Cdt1-mKO2). **(F)** Progression of cells detected in the G1-phase after confinement release, at the beginning of tracking, for 36 hours. **(G)** Progression of cells detected in the G2-phase after confinement release, at the beginning of tracking, for 36 hours. All scale bars represent 5 μm. Statistics in (B) were performed using the Wilcoxon test (* P<0.05, *** P<0.01, *** P<0.001, **** P<0.0001). Statistics in (F) and (G) were performed using the Mann-Whitney-Wilcoxon test (* P<0.05, ** P<0.01, *** P<0.001).

## MATERIALS AND METHODS

### Data and materials availability

All data relevant for the conclusions of this work has been made available through the main text or the supplementary materials. The mass spectrometry proteomics data have been deposited to the ProteomeXchange Consortium via the PRIDE^56^ partner repository with the dataset identifier PXD043524. The ATAC-Seq data have been deposited to the GEO Repository (GSE248846). All code to produce the figures relevant to proteomics analysis in this paper can be found at https://github.com/Skourtis/Rito_Fabio. All other code used in this manuscript can be found at https://github.com/SdelciLab/CINAPS. Relevant supplementary data have been uploaded as Supplementary Data 1 or Supplementary Data 3. For Supplementary Data 2, relevant for cell cycle analysis, please contact authors due to size limitations.

### Cell culture

Unless otherwise mentioned, HeLa (ATCC), U2OS (ATCC), MIA PaCa-2 (ATCC) and MDA-MB-231 (ATCC) cells were maintained in complete Dulbecco’s Modified Eagle Medium (DMEM, Gibco) containing 10% Fetal Bovine Serum (FBS, Gibco) at 37°C and 5% CO_2_. Cell washes were performed with phosphate buffered saline and detachment using Trypsin (Gibco).

### Subcellular fractionation

Subcellular fractionation was performed using a soft lysis buffer (1% CHAPS (cholamidopropyl dimethylammonium 1 propane sulfonate), PBS) and a harsh lysis buffer (0.1% IGEPAL, 20mM Hepes pH 7.5, 300mM NaCl, 0.2mM EDTA). Both buffers contained “complete” proteinase inhibitors (Roche) according to manufacturer’s directions. Suspension or confined cells (using the static confiner) were left in the soft lysis buffer for 5 min, followed by centrifugation at 1 x 10^3^ rcf to obtain the cytoplasmic fraction as the supernatant, and intact nuclei as the pellet. The pellet was then resuspended in the harsh lysis buffer to obtain the nuclear fraction. Fractionation was validated through Western blot.

### Western Blot

Following acquisition and lysis of samples using respective buffers, samples were homogenized using the ICOPLUS Insulin Syringe (Twister Medical, N14080). Samples were quantified and mixed with 4× Laemmli sample buffer (Bio-Rad) and boiled at 95 °C for 5 min. Proteins were separated by SDS– polyacrylamide gel electrophoresis and detected with the following antibodies: Vinculin (Santa Cruz Biotechnology sc-25336), Tubulin (Sigma-Aldrich, T9026), H3 (Cell Signaling Technology 14269) and H3K27me3 (Diagenode, C15410195) antibodies.

### Proteomics

#### Sample preparation for LC-MS/MS analysis

For mass spectrometry analysis, samples were processed using adapted Single-Pot solid-phase-enhanced sample preparation (SP3) methodology (01). Briefly, equal volumes (125 μl containing 6250 µg) of two different kinds of paramagnetic carboxylate modified particles (SpeedBeads 45152105050250 and 65152105050250; GE Healthcare) were mixed, washed three times with 250 µl water and reconstituted to a final concentration of 50 μg/μl with LC-MS grade water (LiChrosolv; MERCK KgaA). Samples (∼100 µL) were filled up to 200 µL with LC-MS grade water and mixed with 2x sample buffer (4% SDS, 100mM HEPES, pH 8.0) to a final concentration of 2% SDS, and proteins were reduced with a final concentration of 10 mM DTT and incubated at 56°C for 1 hour. After cooling down to room temperature, reduced cysteines were alkylated with iodoacetamide at a final concentration of 55 mM for 30 min in the dark. For tryptic digestion, 400 μg of mixed beads were added to reduced and alkylated samples, vortexed gently and incubated for 5 minutes at room temperature. The formed particles-protein complexes were precipitated by addition of acetonitrile to a final concentration of 70% [V/V], mixed briefly before incubating for 18 minutes at room temperature. Particles were then immobilised using a magnetic rack (DynaMag-2 Magnet; Thermo Fisher Scientific) and supernatant was discarded. SDS was removed by washing two times with 200 μl 70% ethanol and one time with 180 μl 100% acetonitrile. After removal of organic solvent, particles were resuspended in 100 μl of 50 mM NH4HCO3 and samples digested by incubating with 2 μg of Trypsin overnight at 37°C. Samples were acidified to a final concentration of 1% Trifluoroacetic acid (Uvasol; MERCK KgaA) prior to immobilising the beads on the magnetic rack. The whole digest was desalted and concentrated using stage tips with two stacked C18 plugs (Empore; MERCK KgaA). Stage tips were activated with three times 100 µl acetonitrile and equilibrated with three times 100 µl of 0.4% formic acid, 2% TFA in water before loading the samples. Salts were cleaned up with 100 µl of 0.1% TFA and peptides were eluted using two times 50 µl 90% acetonitrile, 0.4% formic acid. Finally, eluates were dried in a vacuum concentrator and reconstituted in 10 µl of 0.1% TFA. After solvent removal in a vacuum concentrator, samples were reconstituted in 5% formic acid for LC-MS/MS analysis.

#### Liquid chromatography coupled to tandem mass spectrometry LC-MS/MS

Mass spectrometry was performed on an Orbitrap Fusion Lumos mass spectrometer (Thermo Fisher Scientific, San Jose, CA) coupled to a Dionex U3000 RSLC nano system (Thermo Fisher Scientific, San Jose, CA) via nanoflex source interface. Tryptic peptides were loaded onto a trap column (Acclaim™ PepMap™ 100 C18, 3μm, 5 × 0.3 mm, Fisher Scientific, San Jose, CA) at a flow rate of 10 μL/min using 2% acetonitrile in 0.1% TFA as loading buffer. After loading, the trap column was switched in-line with a 50 cm, 75 µm inner diameter analytical column (packed in-house with ReproSil-Pur 120 C18-AQ, 3 μm, Dr. Maisch, Ammerbuch-Entringen, Germany). Mobile-phase A consisted of 0.4% formic acid in water and mobile-phase B of 0.4% formic acid in a mix of 90% acetonitrile and 10% water. The flow rate was set to 230 nL/min and a 90 min gradient used (4 to 24% solvent B within 82 min, 24 to 36% solvent B within 8 min and, 36 to 100% solvent B within 1 min, 100% solvent B for 6 min before re-equilibrating at 4% solvent B for 18 min). Analysis was performed in a data-dependent acquisition mode. MS^57^ spectra were acquired with a mass range of 375-1650 m/z in the Orbitrap at a resolution of 120,000 (at m/z 200). Automatic gain control (AGC) was set to a target of 2 × 105 and a maximum injection time of 50 ms. Precursor isolation width was set to 1.6 Da and HCD normalised collision energy to 30%. MS2 scans were acquired in the linear ion trap (LIT) in rapid scan mode using higher energy collision-induced dissociation (HCD) and a normalised collision energy of 30%. Intensity threshold for precursor selection was set to 1 × 103 with a quadrupole isolation window of 1.6 Da. Automatic gain control (AGC) was set to a target of 1 × 104 and a maximum injection time of 50 ms. Additional parameters were MIPS enabled for peptide selection, charge state inclusion of 2-6 and dynamic exclusion for selected ions set to 60 seconds. A single lock mass at m/z 445.120024 for MS1 mass correction was employed^58^. XCalibur version 4.3.73.11 and Tune 3.3.2782.28 were used to operate the instrument. The mass spectrometry proteomics data have been deposited to the ProteomeXchange Consortium via the PRIDE^56^ partner repository with the dataset identifier PXD043524.

#### Data analysis

Acquired raw data files were processed using Proteome Discoverer 2.4.1.15 SP1. Database search within PD 2.4 was done using the Sequest HT algorithm and Percolator validation software node (V3.04) to remove false positives with strict filtering at a false discovery rate (FDR) of 1% on PSM, peptide and protein level. Searches were performed with full tryptic digestion against the human SwissProt database V2017.06 including a common contamination list with up to two miscleavage sites. Oxidation (+15.9949Da) of methionine was set as variable modification, whilst carbamidomethylation (+57.0214Da) of cysteine residues was set as fixed modification. Data was searched with mass tolerances of ±10 ppm and 0.6 Da on the precursor and fragment ions, respectively. Results were filtered to include peptide spectrum matches (PSMs) with Sequest HT cross-correlation factor (Xcorr) scores of ≥1 and high peptide confidence. For analysis, chromatin data were normalised using the normalize_vsn and median_normalisation functions from the DEP^59^ and proDA^60^ packages, respectively. The rest of the pipeline was followed according to the DEP package, with the inclusion of impute.mi() function for protein-imputation from the imp4p package^61^. Known subcellular localizations for proteins were obtained from the pRoloc package^62^. For the enrichment, the clusterProfiler R package was used^63^ while CORREP was used for the correlations and clustering^64^. REVIGO^65^ was used to prune semantically similar GO terms.

### Static cell confiner

Cells were confined under PBS-based 4% (w/v) low melting point agarose (Agarose Ultrapure Low Melting Point, Invitrogen) pads, with a thickness of 6 mm (Figure 1 A, top). Three pistons were used to press down on a plastic disk placed on top of the agarose pad to ensure a homogenous pressure distribution. Treatment buffers or reagents were injected through holes in the disk and through the agarose pads, directly onto the cells. After treatment, the agarose confinement was released, and samples processed accordingly. This confinement approach was used to perform protocols that required introduction of buffers or reagents whilst under confinement.

### Dynamic cell confiner

Cells were confined using a pressurised dynamic confiner (4DCell) similar to the previously established planar micro-confinement methods (Figure 1 A, bottom)^66^. To confine cells at 3 µm height, an Si mould for microspacers was produced by photolithography in a clean room (Nanofabrication Laboratory, ICFO) by depositing an SU-8 resin on a silicon wafer. Using the mould, 10-mm confinement coverslips were prepared with polydimethylsiloxane (PDMS). Coverslips and dishes were plasma cleaned, coated with 0.5 mg/ml PLL(20)-g[3,5]-PEG(2) (SuSoS), and equilibrated in respective cell culture media before each experiment. Confinement chambers were assembled using glass-bottom dishes (#1.5, MatTek Corporation) and a pressure generator (LineUp™ Flow EZ™ Fluigent) connected to a suction cup to change the pressure for tuning the confinement heights. Validation of the correct height was performed by measuring the cell height in an X-Z reslice. This confinement approach was used to perform microscopy-based protocols.

### Cell staining and drug treatments

For the visualisation of mitochondria and lysosomes, live cells were incubated using 50 nM MitoTracker Deep Red (Thermo Fisher, M22426) or 75 nM LysoTracker Red DND-99 (Thermo Fisher, L7528), respectively, for 30 min. For the visualisation of late endosomes or peroxisomes, live cells were incubated for 16 hours with respective CellLight BacMam 2.0 probes (Thermo Fisher, Peroxisome: C10604, Late Endosome: C10588). To visualise nuclei, 1 mg/ml DNA-Hoechst was incubated for 5 min. Post incubation, cells were trypsinized and resuspended in complete DMEM.

For imaging experiments using transient expression, live cells were nucleofected using the Lonza Cell Line Nucleofector Kit V (VCA-1003) 48 hours prior to the experiment. Whole cell visualisation was done using pmaxGFP (included in the Lonza kit). Inner nuclear membrane visualisation was done using a pCS2-LAP2β-eGFP plasmid^1^. ER visualisation was done using a pmTurquoise2-KDEL plasmid (Addgene #36204), a gift from Dorus Gadella^67^. Nuclear ATP was visualised using the previously published fluorescence resonance energy transfer based ATeam ATP sensors^24^.

For drug treatments, cells were incubated in media containing Oligomycin A (1 μM, MedChem Express HY-16589), CK666 (100 μM, Tocris #3950), Latrunculin A (500 nM, Sigma L5163), or BAPTA-AM (10 μM, Cayman #15551.5) for 30 min; and SMIFH2 (100 μM, Sigma S4826) for 1 hour. Experiments involving Oligomycin A treatment was carried out in glucose-free DMEM for both treatment and control conditions. During imaging, cells were maintained in media containing the respective drugs at the indicated concentrations.

For immunofluorescence, cells were seeded in a tissue culture treated, black, 96-well optical bottom plates (Perkin Elmer, 6055308) and, following indicated drug treatments, fixed using 4% PFA (Thermo Fisher Scientific, 28908) for 20 min at room temperature. Fixed cells were then permeabilised using 0.25% Triton X-100 in PBS for 15 min at room temperature and blocked using 1% BSA in PBST (PBS+ 0.1% Tween 20) for 30 min at room temperature. α-Tubulin staining was performed using 1:1000 anti-α-Tubulin primary antibody (Sigma Aldrich, T6199) for 1 hour at room temperature. Actin staining was performed using 1:40 Phalloidin (Thermo Fisher Scientific, A12379) together with 1:250 Alexa Fluor 633 secondary antibody (Thermo Fisher Scientific, A-21050) for 1 hour at room temperature. 1:1000 DAPI (Sigma-Aldrich, MBD0015) staining was performed for 5 min at room temperature. All steps were separated by 2-3 PBS washes. Fixed and stained cells were stored in PBS at 4°C until imaged.

### Microscopy

Image acquisition was done using 63x NA 1.4 oil objective and at 0.3 µm spacing between z-slices. All live cell microscopy was performed at 37°C and 5% CO_2_. Confocal fluorescence images were acquired using one of two systems.

The Leica STELLARIS microscope was equipped with 405 nm, 448 nm, 488 nm semiconductor lasers and a white light laser source (Leica Microsystems, Wetzlar, Germany). Fluorescence was detected using HyD detectors in photon counting mode for quantification. Transmission light was collected using a forward PMT. For FRET measurements, a 448 nm laser and a 514 nm white light laser was used for CFP and YFP excitation, respectively, and emitted photons were collected using a HyD detector in photon counting mode.

The Leica TCS SP8 microscope was equipped with 405 nm, 488 nm, 561 nm, and 633 nm semiconductor lasers. Fluorescence was detected using two PMT detectors and/or a HyD detector. For FRET measurements, the CFP was excited using the 458 nm and the YFP with the 514 nm lasers and emitted photons collected using the PMTs. Microscopy on the Leica TCS SP8 and Leica STELLARIS microscopes was performed at the Advanced Light Microscopy Unit, CRG, Barcelona.

Live cell time-lapse imaging was performed using the Operetta High Content Screening System (Perkin Elmer) at 37°C and 5% CO2, equipped with a 20x air high NA objective and 8 LEDs (excitation at 365 nm, 405 nm, 440 nm, 475 nm, 475 nm, 510 nm, 550 nm, 630 nm, 660 nm). Experiments were performed using excitation/emission filters 390-420/430-500 for Turquoise2, 460-490/500-550 for Clover, 530-560/570-650 for mKO2 and 615-645/655-760 for Maroon1.

### Image analysis

Raw image data were analysed and quantified using the FIJI software^68^ or Matlab^69^.

#### Nucleus-associated mitochondria (NAM) quantification

Nuclear Regions of Interests (ROIs) were defined for individual cells based on the nuclear Hoechst signal at every z-plane. For NAM quantification, ROIs were then enlarged by 10% to measure the mitochondria that were present around the nucleus. Statistics and plots were generated based on the mean amount of NAM per cell, quantified as the average of the mean MitoTracker intensity per z-plane.

#### Nucleus-mitochondria co-localization analysis

Representative cell line profile (Figure 2 E) was measured along lines of defined distances with a 10-pixel thickness on a sum projection image. i) Line was drawn from the centre of the nucleus to the cell periphery; ii) Line graphs of signal intensities were depicted as standardised signal intensity using Matlab.

#### Organelle co-localization analysis

Signal intensities were measured along lines of defined distances with a 10-pixel thickness. i) NAM was evaluated using lines from the centre of the nucleus to the cell periphery; ii) NAM-ER, NAM-late endosome, NAM-lysosome and NAM-peroxisome general co-localization (wide zoom, Figure 3 E, Figure S3 B, E, H) was performed with 3 measurements per cell, for 5 cells, using lines spanning the longest nuclear diameter. (iii) Detailed NAM-ER, NAM-late endosome, NAM-lysosome, and NAM-peroxisome localization (zoom in, Figure 3 F, Figure S3 C, F, I) was analysed using lines of length 1.5 μm starting at a point of high MitoTracker signal. Line graphs of signal intensities were depicted as mean standardised signal intensity along with their 95% confidence interval. This analysis was performed using a generalized additive model available within the ggplot package^70^.

#### Kymograph and micro-dispersion analysis

Line profiles of ER signal across the longest diameter of the cell was measured for 70 seconds. Kymographs are line profiles (x-axis) across time (y-axis). Signal dispersion was depicted as mean standard deviation across standardised distances along with their 95% confidence interval. This analysis was performed using a generalized additive model available within the ggplot package^70^. Results are represented as mean with 95% confidence intervals.

#### FRET analysis

ATP levels were analysed using an adaptation of previously published protocols^71^. Briefly, a ROI per z-stack was selected for every image. The ratio YFP/CFP was calculated by dividing the FRET image (Ex CFP/ Em YFP) by the CFP image (Ex CFP/ Em CFP). Results are represented as the mean intensity per cell.

#### Pixel binning and coefficient of variation

3 central planes of nuclei were selected and used to produce maximum projection images. DAPI signal (Ex 405/ Em DAPI) was quantified for mean signal intensity, and signal standard deviation. The coefficient of variation was calculated as the ratio of signal standard deviation to mean signal intensity for each nucleus. This process was repeated for the same images using pixel binnings at 1x1 = 1px; 2x2 = 4px, 3x3 = 9px, 4x4 = 16px, 5x5 = 25px, 6x6 = 36px; 7x7 = 49px; and 8x8 = 64px. Results are represented as mean coefficient of variation with 95% confidence intervals.

### ATAC-sequencing sample preparation

HeLa cells were treated with or without Oligomycin A (1 µM) for 30 min in glucose-free complete media. Cells were then trypsinized and either left in suspension (controls) or confined using a hybrid confiner. Briefly, cells were confined in glass-bottom 6-well plates (adapted from 4D Cell, CSOW 610) under coverslips with 3µm height microspacers. Both, coverslips, and the glass-bottom plate were plasma-cleaned and treated with 0.5 mg/ml PLL(20)-g[3,5]-PEG(2) (Susos) to prevent attachment. Following an acute 15-minute confinement, 0.5x10^5^ cells for two biological replicates, for each treated condition, were collected and treated with transposase Tn5 (Nextera Tn5 Transposase; Illumina Cat #FC-121-1030). DNA was purified using AMPure XP beads to remove large fragments (0.5 × beads; >1kb) and small fragments (1.5 × beads; < 100bp). Samples were then amplified using NEBNext high-Fidelity 2x PCR Master Mix (New England Labs #M0541) with primers containing a barcode to generate the libraries, as previously described^72^. Each condition was amplified using a combination of the P7 forward primer and a P5 reverse primers containing multiplexing adaptors (Table 1) and sequenced accordingly to allow for segregation of reads. The number of cycles of library amplification was calculated as previously described^73^. DNA was purified using a MinElute PCR Purification Kit (Qiagen), and samples were sequenced using an Illumina NextSeq2000 at the CRG Genomics Core Facility.

**Table 1.** : ATAC-seq library preparation multiplexing adaptors.

| Replication/Treatment/Confinement | Sample name | Multiplex Index P7 | Multiplex Index P5 |
| --- | --- | --- | --- |
| Replicate 1 Untreated Suspension | n4_UU | TAAGGCGA | TAGATCGC |
| Replicate 1 Untreated Confined | n4_CU | CGTACTAG | CTCTCTAT |
| Replicate 1 Treated Suspension | n4_UT | AGGCAGAA | TATCCTCT |
| Replicate 1 Treated Confined | n4_CT | TCCTGAGC | AGAGTAGA |
| Replicate 2 Untreated Suspension | n5_UU | GGACTCCT | GTAAGGAG |
| Replicate 2 Untreated Confined | n5_CU | TAGGCATG | ACTGCATA |
| Replicate 2 Treated Suspension | n5_UT | CTCTCTAC | AAGGAGTA |
| Replicate 2 Treated Confined | n5_CT | CGAGGCTG | CTAAGCCT |

### ATAC-sequencing analysis

50 base-pairs paired-end reads, were subjected to alignment against the hg19 human genome using Bowtie2 aligner (v 2.5.1)^74^ after removal of adaptors with TrimGalore (v. 0.6.10). The alignment was carried out with default parameters, except for the utilization of the “very-sensitive 2000” option. To ensure the quality of aligned reads, those with a quality score below 30 were filtered out using Samtools 1.16.1^75^. Mitochondrial reads were excluded from further analysis followed by removal of duplicated reads by using Picard (v 2.23.8). The alignment of reads was adjusted to account for Tn5 transposase insertions with deepTools (v 3.5.1)^76^. Peaks were identified using MACS2 (v 2.2.8) with false discovery rate (FDR) threshold of less than 0.01. Regions designated as blacklisted were filtered out from further analysis (blacklist v2). Peak intersection between replicates and union between conditions was done using the GenomicRanges package in R (V4.2). Differentially accessible (DA) regions were quantified utilizing the csaw R package^77^ with Trimmed Mean of M-values (TMM) normalization on binned counts followed by edgeR^78^ for DA analysis, employing a significance threshold of log-fold change > 0.5 and FDR < 0.1.

### Cell cycle analysis

A previously established stable U2OS cell line with a FUCCI system was used^40^. Briefly, our FUCCI system is an adaptation of FUCCI4^39^, showing 3 cell cycle regulated fusion proteins: Clover-Geminin, SLBP-Turquoise2 and Cdt1-mKO2. Fluorescence from tracked live cells was measured using the Operetta High Content Screening System (Perkin Elmer), using 20x magnification (see ‘Microscopy’ methods for more details). Control or treated cells were confined for a duration of 15 min, following which they were released and seeded in cell culture plates. Fluorescence of FUCCI probes were then tracked using live cell imaging for 36 hours. Quantification of cell cycle was based on fluorescence from the FUCCI proteins and was performed using a custom R script. Briefly, fluorescence values were standardised between 0 and 1, and cell cycle status visualised as XYZ plots (x axis = SLBP-Turquoise2, y axis = Clover-Geminin, z axis plotted as colour = Cdt1-mKO2). Cell cycle phase gates – G1, S, G2 and M – were assigned based on their distribution (Figure 5 E). Further analysis to calculate percentage population abundance in various phases, cell cycle progression through various phases, and time spent in a specific phase were all based on the above-mentioned gating strategy. Relative cell proliferation was analysed by normalising total cell count to that of the first time point.

### DNA damage

DNA damage foci were quantified using a stable U2OS cell line expressing an mMaroon1-53BP1trunc reporter. Briefly, the reporter cell line was created by replacing the mApple fluorophore in the Apple-53BP1trunc Addgene vector #69531, a gift from Ralph Weissleder^37^, with an mMaroon1 fluorophore derived from the Addgene vector #83842, a gift from Michael Lin^39^. Using this vector, a stable U2OS cell line was created using lentivirus transduction. Cells were either treated or left untreated with Oligomycin A (1 μM) as described above. Confinement was performed using the dynamic confiner and confocal microscopy to quantify foci during confinement. Quantification was done using Fiji^68^ and was based on maximum projections to detect individual foci. To quantify foci post confinement release, confinement was released, and cells were seeded in 96-well plates under normal growth conditions and fluorescence from tracked live cells was measured using the Operetta High Content Screening System (Perkin Elmer).

